# Dearth of smoking-induced mutations in NSRO-driven non-small-cell lung cancer despite smoking exposure

**DOI:** 10.1101/2023.07.04.547310

**Authors:** Chen-Yang Huang, Nanhai Jiang, Meixin Shen, Gillianne Lai, Aaron C. Tan, Amit Jain, Stephanie P. Saw, Mei-Kim Ang, Quan Sing Ng, Darren Wan-Teck Lim, Ravindran Kanesvaran, Eng-Huat Tan, Wan Ling Tan, Boon-Hean Ong, Kevin L. Chua, Devanand Anantham, Angela Takano, Tony K.H. Lim, Wai Leong Tam, Ngak Leng Sim, Anders J. Skanderup, Daniel S.W. Tan, Steven G. Rozen

## Abstract

Non-small cell lung cancers (NSCLCs) in non-smokers are mostly driven by mutations in the oncogenes *EGFR, ERBB2,* and *MET*, and fusions involving *ALK* and *RET*. We term these “non-smoking-related oncogenes” (NSROs). In addition to occurring in non-smokers, NSRO-driven tumors also occur in smokers, and the clonal architecture and genomic landscape of these tumors remain unknown. We investigated genomic and transcriptomic alterations in 173 tumor sectors from 48 patients with NSRO-driven or typical-smoking NSCLCs. NSRO-driven NSCLCs in smokers and non-smokers have similar genomic landscapes. Surprisingly, even in patients with prominent smoking histories, the mutational signature caused by tobacco smoking was essentially absent in NSRO-driven NSCLCs. However, NSRO-driven NSCLCs in smokers had higher transcriptomic activities related to regulation of the cell cycle, suggesting that smoking still affects tumor phenotype independently of genomic alterations.

**Statement of significance:** This study highlights the lack of genomic scars caused by smoking in NSCLCs driven by non-smoking-related oncogenes regardless of smoking history. The impact of smoking on these tumors is mainly non-genomic. The transcriptomic features of NSCLCs associated with smoking may help in the development of therapeutic approaches.

## Introduction

Lung cancer remains the most lethal cancer worldwide and causes more than 1.8 million deaths annually, even though the worldwide prevalence of tobacco smoking is decreasing (1, 2). However, in East Asia, lung cancer among non-smokers is increasing, has become an emerging health problem, and accounts for more than half of cases in Taiwan and Singapore (3, 4). Most of these are non-small-cell lung cancers (NSCLCs) driven by a specific set of oncogenic mutations, usually activating mutations in the oncogenes *EGFR*, *ERBB2*, or *MET*, or rearrangements involving *ALK* or *RET* (5-9). Here, we refer to these genes as “non-smoking-related oncogenes” (NSROs). Including both smokers and non-smokers, NSRO-driven NSCLCs constitute approximately half of all East-Asian lung cancer (10-13). These tumors tend to have similar clinical trajectories and favorable responses to tyrosine kinase inhibitors (TKIs) (14-21). In contrast, tumors that occur in smokers and that have activating mutations in the *KRAS* or *BRAF* genes or that lack mutations in known oncogenes altogether constitute a group that we term “typical-smoking NSCLCs”. Typical-smoking NSCLCs are more common in populations of European descent than in East-Asian populations. They often have high mutational burdens, sometimes respond well to immune checkpoint inhibitors, and have many somatic mutations caused by tobacco smoking (22, 23). In contrast, NSRO-driven NSCLCs tend to have lower mutational burdens and complex genomic architectures (24, 25).

While NSRO-driven NSCLCs have been most studied in non-smokers, in East Asia, approximately 20% to 40% of patients with these cancers have histories of tobacco smoking (4, 5, 26-28). Furthermore, among patients with *EGFR*-mutated NSCLC treated with TKIs, smokers have worse prognoses than non-smokers (29-31). However, the impact of tobacco smoking on clonal architecture, somatic genomic alterations, and transcriptomic phenotypes in NSRO-driven NSCLCs remains largely unknown. By means of an integrated study of genomic and transcriptomic landscapes, clonal architecture, and intra-tumor heterogeneity, we investigated similarities and differences between (i) NSRO-driven NSCLCs in non-smokers (ii) NSRO-driven NSCLCs in smokers and (iii) typical-smoking NSCLCs.

## Results

### Characteristics of the study cohort

We performed multi-region exome sequencing in a total of 173 sectors from 48 NSCLCs with clinical and histopathological characteristics shown in Supplementary Tables S1 and S2. Overall, we identified 6,251 single-nucleotide variants (SNVs) and 314 small indels (insertions or deletions) affecting the exons of 4,738 genes and the splicing junctions of 177 genes. We also performed RNA sequencing on 103 of the 173 sectors from 32 out of the 48 tumors (Supplementary Tables S1, S2).

Consistent with previous studies in East-Asian NSCLC, *EGFR* and *TP53* were the most frequently mutated genes (56%, 27 of 48, and 46%, 22 of 48, respectively) (9, 10). In addition to the SNVs and indels, we also studied gene fusions using RNA sequencing data. Of the putative transcript fusions detected, three were known oncogenic variants: *EML4*-*ALK*, *KLC1*-*ALK*, and *PARG*-*BMS1* (Supplementary Table S3), and the two *ALK* fusions were confirmed by fluorescence in situ hybridization (32-34). As expected, mutations in *EGFR*, *MET*, *ERBB2*, and *KRAS* and *ALK* fusions did not co-occur, suggesting that these were key initiating events (35).

All *EGFR*, *ERBB2,* and *MET* mutations were truncal (occurred in every sector) and were clonal in every sector (Supplementary Table S4). Compared to mutations in non-driver genes, mutations in *EGFR* were statistically more likely to be truncal (Supplementary Table S5). These findings underscore the impact of these oncogene mutations on the clonal architecture of NSCLCs. In addition, mutations in *KRAS* were truncal in every tumor, but were subclonal in single sectors in 2 out of 7 of the tumors. A similar pattern was reported in another multi-regional study (36).

### Three categories of NSCLC and their genomic characteristics

For the purposes of this study, we first defined NSRO-driven tumor as those with any of the mutations or other genetic alterations detailed in Supplementary Table S6, based on the incidences of these alterations reported in the literature (27, 28). The NSRO mutations observed in this study were activating mutations in *EGFR* exons 18 through 21, *ALK* fusions, insertions in *ERBB2* exon 20, and skipping of *MET* exon 14.

We then focused on the following three groups: (i) NSRO-driven tumors in non-smokers (n = 23, 48%), (ii) NSRO-driven tumors in smokers (n = 12, 25%), and (iii) typical-smoking tumors (n = 11, 23%). In addition, there were 2 tumors without NSRO-mutations in non-smokers. This is typical of East-Asian populations, in which tumors without NSROs are rare in non-smokers (7, 8, 28). The proportion of non-smokers without NSROs in this study (8%, 2 out of 25) was lower than those reported in population of European descent (e.g., 51%, 96 out of 189, p = 0.05 by Fisher’s two-sided exact test) (37). Fig. 1A shows the genomic landscape of the three groups.

**Fig. 1.**
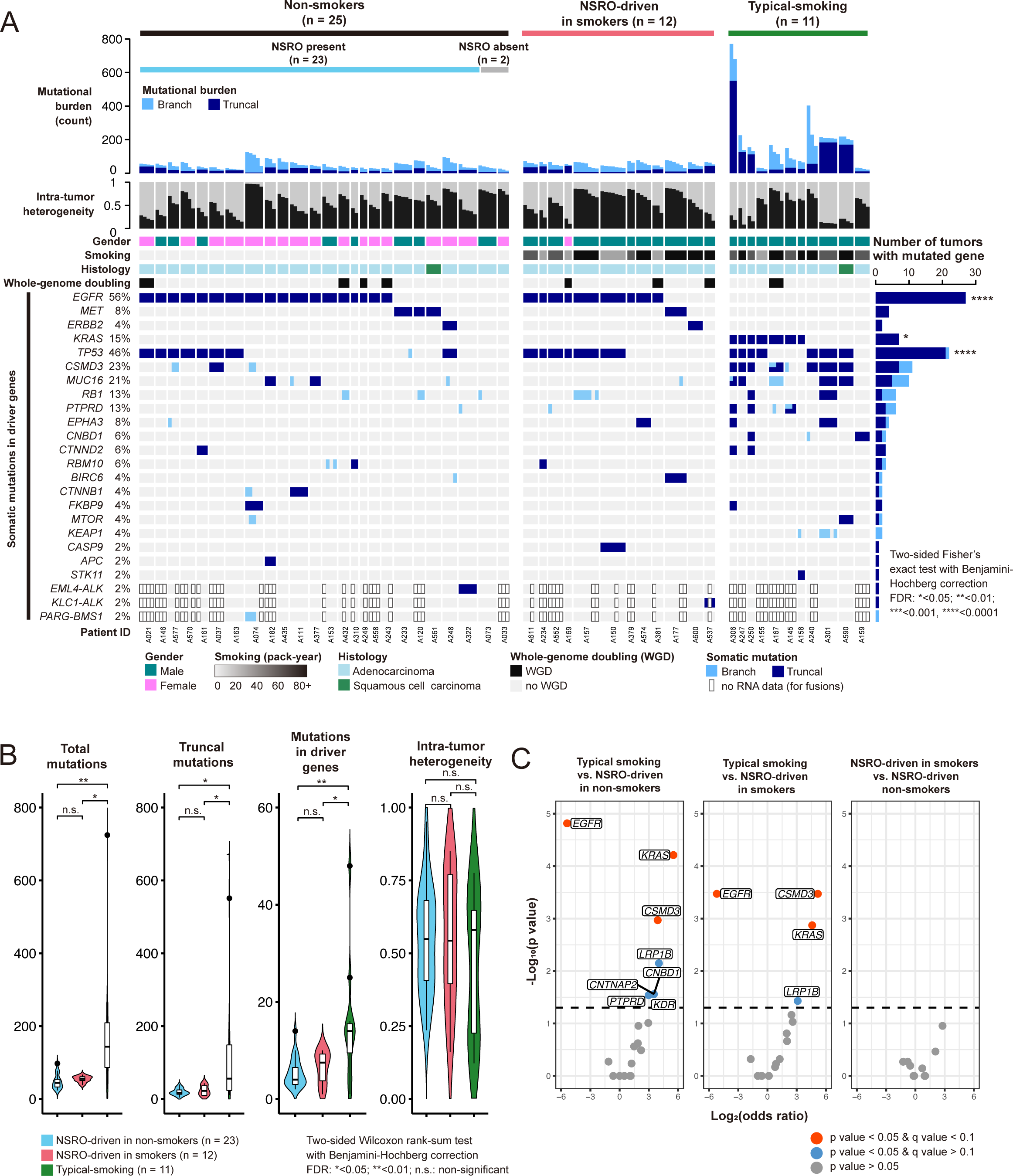
(A) Overview of genomic alterations in tumors and tumor sectors. (B) Counts of total mutations, truncal mutations, numbers of mutations in driver genes, and levels of intra-tumor heterogeneity in the three groups. We used the list of driver genes from COSMIC (https://cancer.sanger.ac.uk/census). (C) Enrichment for mutations in driver genes in pair-wise comparisons among the three groups.

Table 1 summarizes clinical characteristics and NSRO mutations in the three groups. Notably, smoking exposure was similar between NSRO-driven tumors in smokers (median 34.5 pack-years, range 0.5-99) and typical-smoking tumors (median 38 pack-years, range 2-168, Wilcoxon rank-sum test, p value = 0.5792).

**Table 1.**
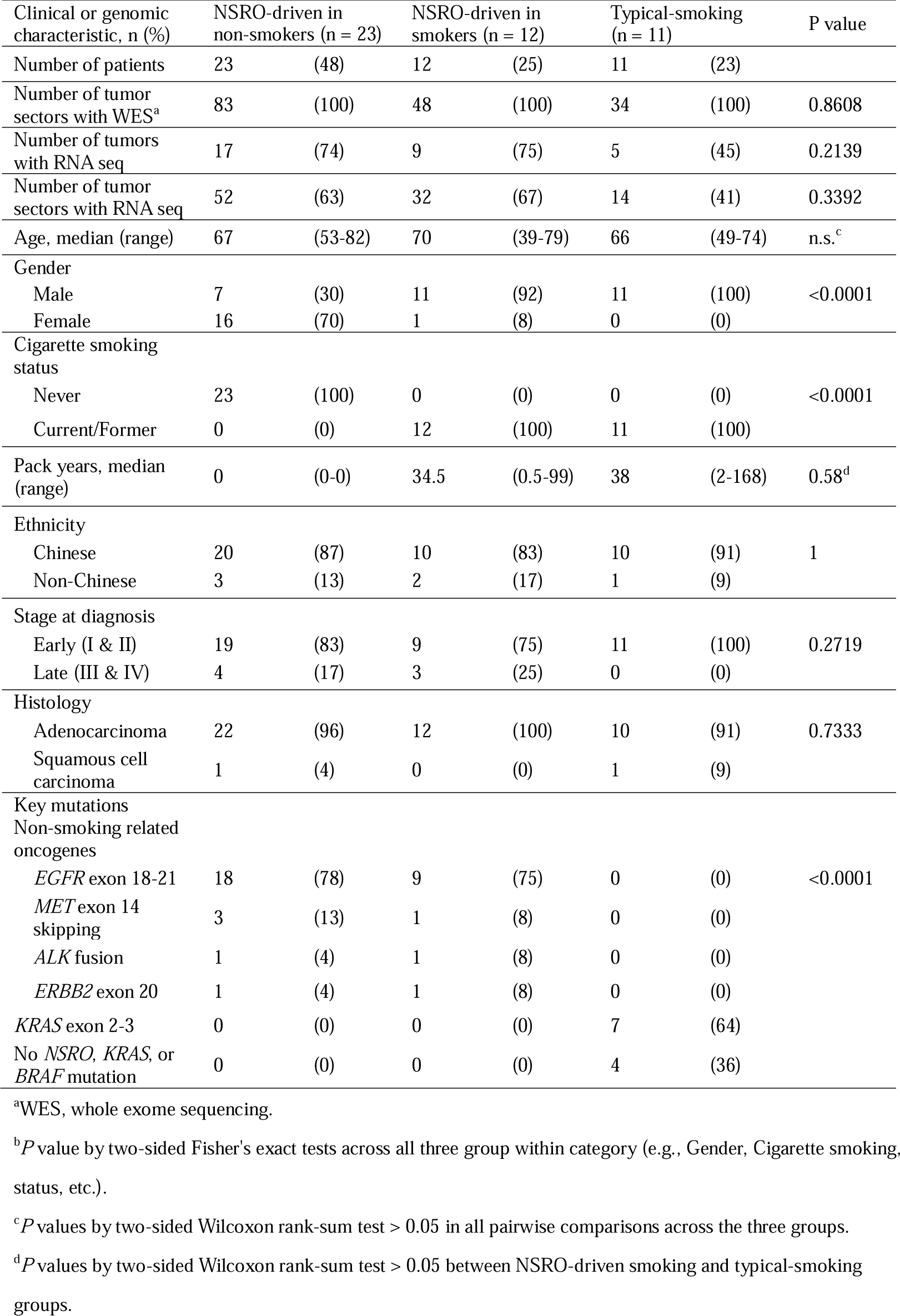
Baseline clinical and genomic characteristics of NSRO-driven NSCLCs in non-smokers, NSRO-driven NSCLCs in smokers, and typical-smoking NSCLCs.

### NSRO-driven NSCLC with and without smoking histories have similar genomic architectures

Overall, NSRO-driven tumors in smokers and non-smokers had similar genomic architectures, including tumor mutational burdens (TMB), number of truncal mutations, and number of mutations in driver genes. In contrast, compared to NSRO-driven tumors in smokers, typical-smoking tumors had much higher TMBs (median 144 vs. 55.5, p = 0.017, two-sided Wilcoxon rank-sum test), more truncal mutations (median 56 vs. 22.5, p = 0.031), and more mutations in all driver genes (including NSROs, median 14 vs. 7.5, p = 0.002, Fig. 1B, Supplementary Data and Code). Intra-tumoral heterogeneity (ITH, defined as the mean ratio of the numbers of branch mutations to the total number of mutations, see Methods) was similar across the three groups (medians 0.549, 0.543, and 0.580, for NSRO-driven non-smokers, NSRO-driven NSCLCs in smokers, and typical-smoking NSCLCs, respectively, Fig. 1B). However, “coconut-tree” phylogenies, characterized by a combination of high TMB (> 100) and low ITH (< 0.5), occurred exclusively among the typical-smoking tumors (5 out of 11, Fig. 2). Supplementary Fig. S1 details phylogenetic trees of all tumors across the three groups.

**Fig. 2.**
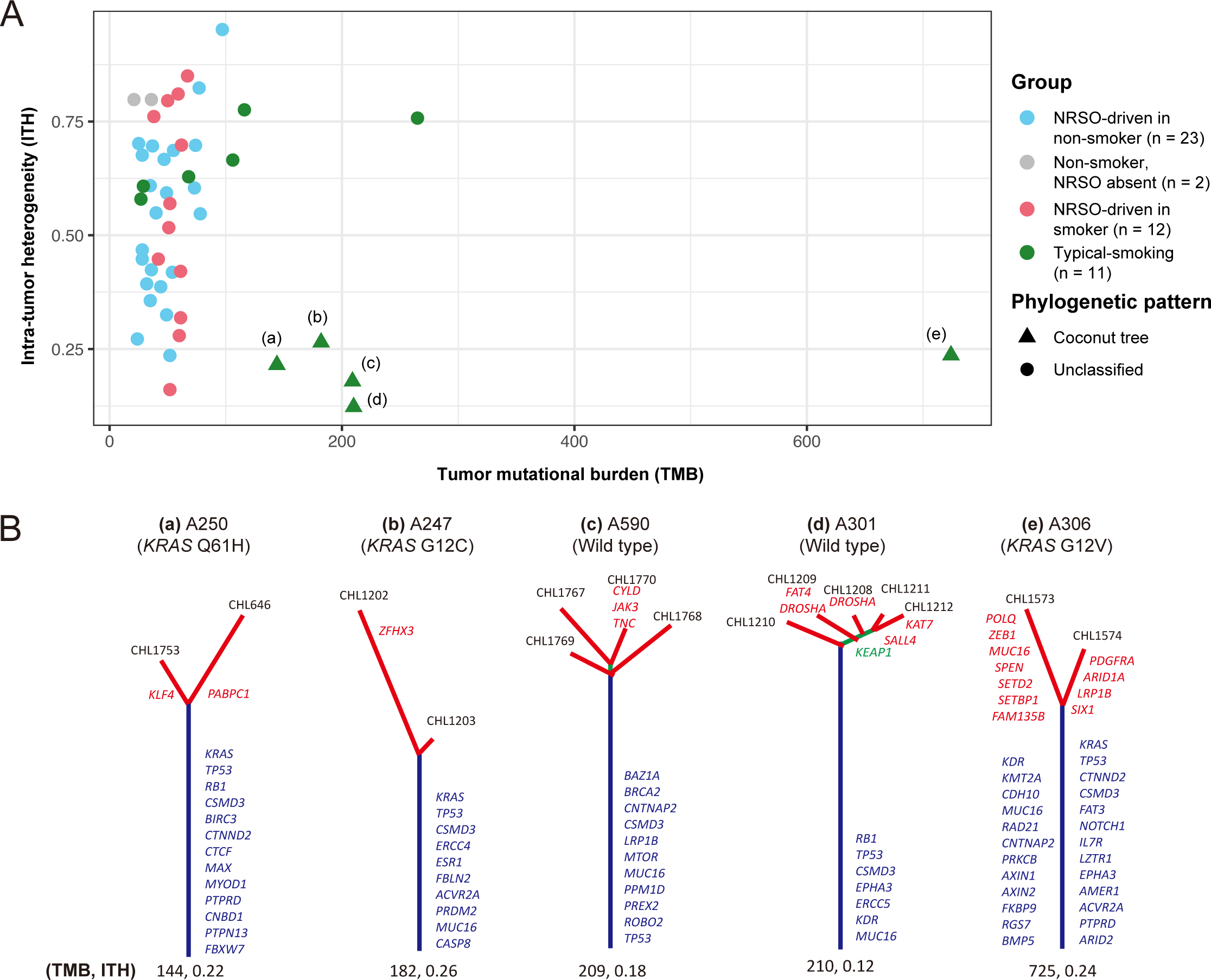
(A) Intra-tumor heterogeneity (ITH) versus tumor mutation burden (TMB) for each tumor. Five tumors with “coconut-tree” phylogenies are labeled (a) through (e) and the corresponding phylogenies are in panel (B). These phylogenies occurred only in the typical-smoking-related group.

In addition to the oncogenes used to categorize the three groups of tumors, *CSMD3* mutations were statistically more common in typical-smoking tumors (Fig. 1C, left and middle). In comparing NSRO-driven NSCLCs in smokers versus NSRO-driven NSCLCs in non-smokers, there was no significant difference in the prevalence of mutations in COSMIC driver genes (including NSROs, Fig. 1C, right, https://cancer.sanger.ac.uk/census).

Previous studies reported that whole-genome doubling and chromosomal instability are common features of NSCLC (24, 25, 38, 39). In the present study, we found no significant difference across the three groups in tumor ploidy, proportions of tumors with whole-genome doubling, and chromosomal instability (Supplementary Fig. S2, Supplementary Table S2). Supplementary Fig. S3 provides details of somatic copy number alterations for all groups.

We also noted that gender distribution differed strongly across the three groups. Among non-smokers with NSRO-driven tumors, only 30% (7 of 23) were male, whereas among the NSRO-driven smoking and typical-smoking groups, 92% and 100% were male, respectively (p = 0.0024 and 0.0001 by two-sided Fisher’s exact tests compared to the NSRO-driven non-smoking group). We analyzed genomic landscapes in NSRO-driven tumors by gender and found no significant differences (Supplementary Fig. S4).

### Mutational signatures of NSRO-driven NSCLCs in smokers

Next, we investigated the impact of smoking on the mutational landscape across three groups. Overall, the median number of single-base substitution (SBS) per sector was 173 (range 47 to 2472). We used a signature presence test followed by signature attribution for each tumor sector to detect the mutational signature SBS4, which is caused by tobacco smoking in lung cancers (22, 40). Fig. 3A shows the activities of mutational signatures of the three groups.

**Fig. 3.**
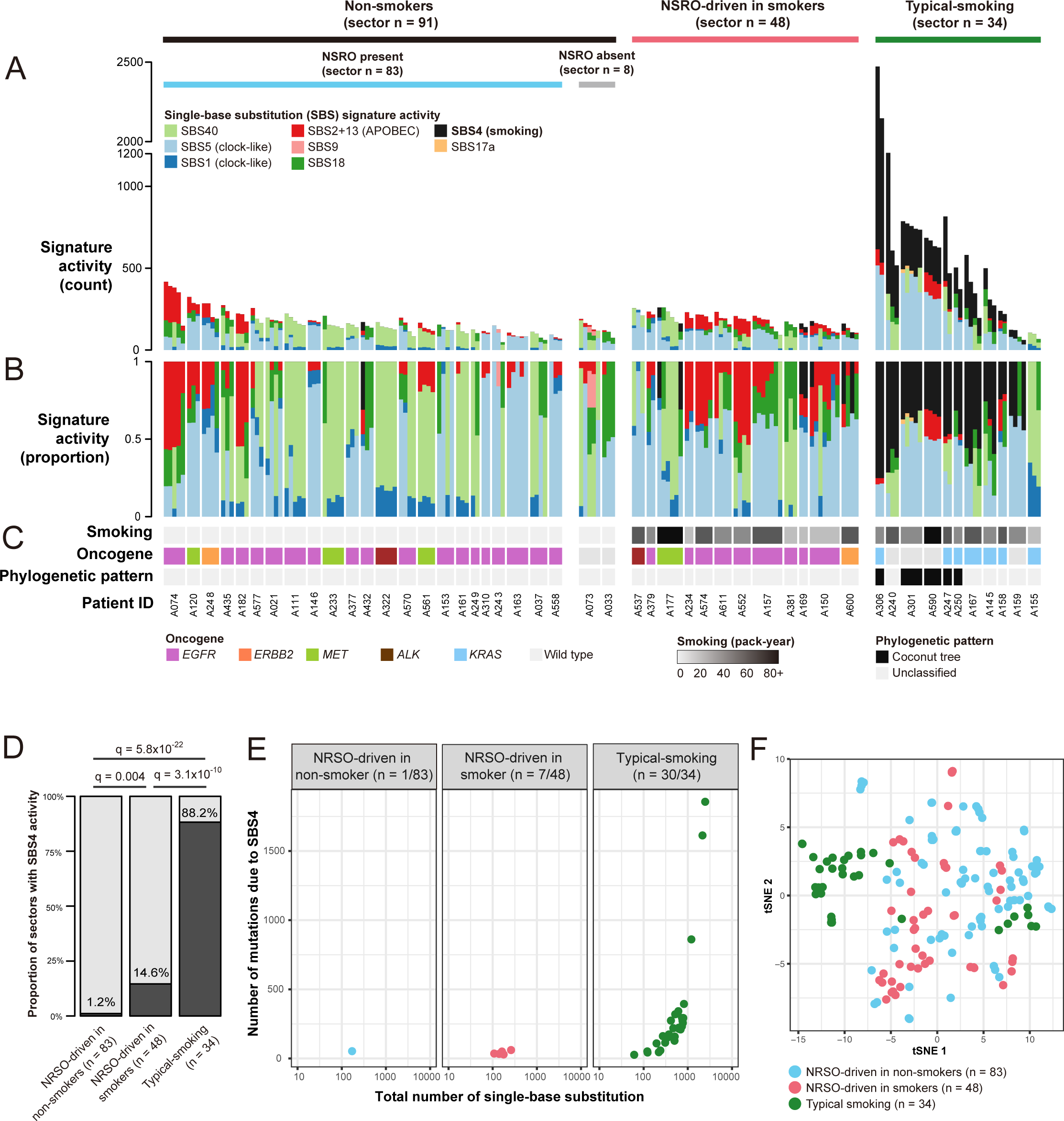
Single-base substitution (SBS) mutational signatures. (A, B) Mutational-signature activities in the three groups by absolute mutation counts (A) and by proportions (B). Colors indicate various mutational signatures (e.g., SBS1, SBS5, etc.), as indicated by the legend above. (C) Smoking history, key mutated genes, and whether the tumor has a coconut tree pattern. (D) Proportions of tumor sectors with SBS4 (caused by tobacco smoking). (E) Counts of mutations due to SBS4 in tumor sectors that have SBS4 mutations. (F) tSNE (t-distributed stochastic neighbor embedding) dimension reduction based on the mutational spectra. For information on 10the mutational signatures, see COSMIC (https://cancer.sanger.ac.uk/signatures/).

We detected SBS4 in 30 of 34 (88%) sectors in typical-smoking tumors. Surprisingly, however, SBS4 was found in only 7 of 48 (15%) sectors in NSRO-driven tumors in smokers, significantly less than sectors in typical-smoking tumors despite similar exposures to tobacco smoking (two-sided Fisher’s exact test, p < 2.1×10^-11^, Fig. 3D). For tumor sectors with SBS4 activity, the median number of mutations attributed to SBS4 was also significantly higher in typical-smoking tumors than in NSRO-driven tumors in smokers (216 vs. 53, two-sided Wilcoxon rank-sum test, p < 9×10^-5^, Fig. 3E). T-distributed stochastic neighbor embedding (tSNE) based on SBS spectra identified different mutational patterns in typical-smoking tumors compared to the other two groups (Fig. 3F, Supplementary Table S7 and S8).

To confirm the paucity of SBS4 activity in NSRO-driven NSCLCs, we applied the same tumor classification and signature assignment algorithm to two large, previously reported cohorts of NSCLC (10, 41). Supplementary Table S9 provides clinical information, including smoking history, oncogenes and their mutations, and signature activities for patients in these cohorts. The SBS4 signature was found in only 38% (8 of 21) and 29% (9 of 31) of NSRO-driven NSCLCs in smokers, significantly fewer than in typical-smoking tumors (90% and 78%, odds ratio of 0.07 and 0.12, two-sided Fisher’s exact tests, p values of 1.1×10^-7^ and 4.5×10^-6^, respectively, Supplementary Figs. S5, S6). Thus, all the genomic data indicates that NSRO-driven NSCLCs, whether in smokers or non-smokers, have origins and oncogenic histories distinct from those of typical-smoking NSCLCs.

Previous studies of NSCLC suggested that mutations caused by smoking and *APOBEC* activities dominate at different stages of cancer evolution (24, 42). For smoking mutagenesis, in the current study, SBS4 contributed similar activities in trunks and branches in typical-smoking tumors, suggesting ongoing exposure to tobacco smoke during cancer development (Supplementary Fig. S7). Analysis of SBS4 was not meaningful for the other two groups of NSCLCs, because they had almost no SBS4 mutations. For *APOBEC*, consistent with previous studies, there were more mutations in branches than in trunks across the entire data set (Supplementary Fig. S7). Unexpectedly, we found that mutations due to reactive oxygen species (ROS, SBS18) were significantly higher in the branches than in the trunks for every group of tumors (all q values < 0.018 by two-sided Wilcoxon paired rank-sum tests with Benjamini-Hochberg correction, Supplementary Fig. S7). Thus, ROS might contribute to NSCLC evolution.

### Transcriptomic features of NSRO-driven smoking tumors

The similarity of genomic landscapes between NSRO-driven tumors with and without smoking histories was surprising because clinical studies have shown that smoking indicates poor prognosis in NSRO-driven NSCLC (29-31). Therefore, we investigated whether transcriptomic features may account for this clinical observation. To this end, we profiled the transcriptomes of 103 of the 173 sectors from 32 out of the 48 tumors (Table 1 and Supplementary Table S2). UMAP dimension reduction did not reveal a separation between NSRO-driven NSCLCs in smokers versus non-smokers in the first 2 dimensions (Supplementary Fig. S8A). Indeed, the primary separation seems to reflect membership in the terminal-respiratory-unit (TRU) expression subtype. There is a trend for association of the TRU subtype with NSRO-driven tumors in both non-smokers and smokers compared to typical-smoking tumors (Supplementary Fig. S8B).

To further explore the transcriptomic activities associated with tobacco smoking, we investigated differential expression pathway activity between non-smokers (57 sectors from 18 tumors) and smokers (46 sectors from 14 tumors, including both typical-smoking tumors and NSRO-driven tumors in smokers). We examined activities of 1,259 pathways from the Reactome Database (43) (Fig. 4, Supplementary Fig. S9, Supplementary Table S10). In this comparison, tumors in non-smokers had higher activities in pathways related to NOTCH signaling and to metabolism and lower activities in pathways related to cell cycle regulation and mitotic exit (Fig. 4). These differences in pathway activity were also evident in a comparison of NSRO-driven tumors in smokers versus in non-smokers (i.e., after excluding typical-smoking tumors and NSRO-negative tumors, Supplementary Table S10). These observations underscore the phenotypic differences associated with tobacco smoking in some transcriptomic pathways independent of genomic alterations.

**Fig. 4.**
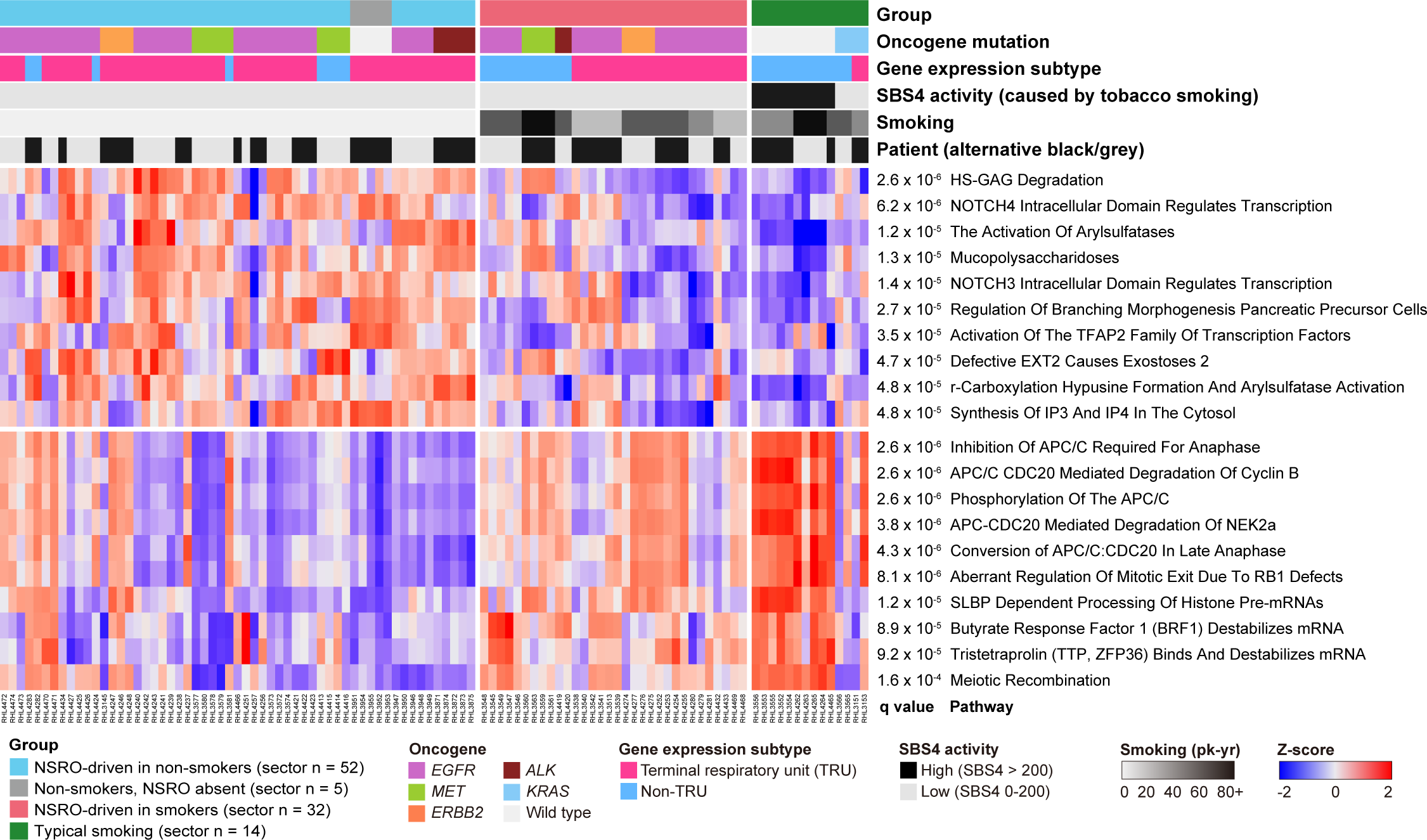
Heatmap of activities of the top 10 pathways up- and down-regulated in all non-smokers compared to all smokers. Each column is a tumor sector, and sectors are grouped by patient as shown in the row labeled “Patient”. Z-scores are of pathway activity. The Benjamini-Hochberg false discovery rates (*q* values) of differential pathway activity were based on *p*-values calculated using limma (84). Supplementary Fig. S9 and Supplementary Tables S10 and S11 provide details.

It has been proposed that smoking-associated lung cancers are associated with the immune repertoires of tumor microenvironments (TME) that would confer better responses to immunotherapy (44-47). TMEs with infiltration of cytotoxic T cells and expression of proinflammatory cytokine genes are often called “immune-hot”, and have higher tendencies to respond to immune checkpoint inhibitors (48, 49). In this study, we investigated the TME of NSRO-driven NSCLCs in smokers versus the other two groups. To define immune-hot TME, we performed hierarchical clustering of sectors based on the transcript levels of T-cell inflammation and immune checkpoint genes (Supplementary Fig. S10A) (50-52). However, there was no strong evidence for enrichment for immune-hot TMEs in NSRO-driven NSCLCs in smokers (3 out of 9 tumors, 33%) and typical-smoking-related NSCLCs (1 out of 5 tumors, 20%) compared to NSRO-driven tumors in non-smokers (5 out of 17 tumors, 29%, p = 0.7332 by Fisher’s exact test across the three groups, Supplementary Fig. S10B).

## Discussion

To our knowledge, this is the first integrated genomic study to directly compare NSRO-driven NSCLCs in non-smokers, NSRO-driven NSCLCs in smokers, and typical-smoking NSCLCs using multi-region exome and RNA sequencing. We found that tobacco smoking had almost no influence on the genomic features and clonal architectures of *EGFR*-mutated and other NSRO-driven NSCLCs. Despite prominent smoking histories, NSRO-driven tumors in smokers were similar to those in non-smokers in terms of mutational burden, intra-tumor heterogeneity, and mutational signature activity. In contrast, compared to both NSRO-driven groups, typical-smoking tumors showed higher TMBs and more mutations in driver genes. Furthermore, “coconut-tree” phylogenies, which are defined by a combination of high TMB (> 100) and low ITH (< 0.5), occurred in nearly half of the typical-smoking NSCLCs but were absent from NSRO-driven NSCLC.

As noted in the Results section, gender distribution differed significantly across the three groups. Across all groups, tobacco smoking was more prevalent among males than females (Table 1). This male preponderance reflects the extreme gender imbalance of smoking in East Asia. For example, in the population we studied, 6.8% of women are smokers compared to 20.6% of men (2, 53, 54). Previously, NSRO-driven NSCLC was sometimes viewed as a disease of non-smokers, often women. This view may have been partly driven by this gender imbalance, due to which NSRO-driven tumors were particularly noticeable among women, since they were usually non-smokers. The current study confirms that NSRO-driven NSCLC occurs in both smokers and non-smokers and in both sexes, and it shows that genomic features are similar in both smokers and non-smokers and in both sexes (Supplementary Fig. S4). Because of the strong differences in smoking rates between women and men in the study population, it is not possible to disentangle the effects of gender from the effects of smoking. Nevertheless, we note that available evidence suggests that NSRO-driven tumors are more common among women. Notably, in data from a previous study (55), in both non-smokers and smokers, *EGFR*-mutated NSCLC was more common among women: for non-smokers, odds-ratio 1.38 (p < 8× 10^-9^); and for smokers, odds ratio 1.40 (p < 0.006), analyses by two-sided Fisher’s exact tests.

While it has been recently reported that NSCLCs that occur in smokers but that lack SBS4 are enriched for mutations in *EGFR* and other NSROs (36, 56), here we have shown that the paucity of mutations due to tobacco smoking is a nearly-universal characteristic of NSRO-driven tumors in smokers. We further confirmed this finding in two large cohorts of patients with lung adenocarcinomas (10, 41). It is unclear why NSRO-driven tumors rarely acquire mutations caused by smoking, while typical-smoking tumors with similar exposures have abundant smoking mutations. Possibly, as suggested by some studies, the cell of origin of NSRO-driven NSCLC may be different from that of typical-smoking NSCLC (57-61). Thus, one possibility is that NSRO-driven tumors are less prone to mutation because their cells of origin are less exposed to tobacco smoke or have more effective DNA damage repair.

Although NSRO-driven tumors in non-smokers and in smokers have similar clonal architectures and genomic features, they differ in their transcriptomic pathway activities, especially those related to the cell cycle and mitotic exit. Consistent with previous studies (62, 63), for these pathway activities, NSRO-driven tumors in smokers are more similar to typical-smoking tumors than to NSRO-driven tumors in non-smokers (Fig. 4, Supplementary Fig. S9). Despite the lack of somatic mutations caused by smoking, it is still possible that chronic tobacco exposure causes epigenomic changes to bronchial epithelial cells, leading to a carcinogenic phenotype that is independent of genomic alterations (64). Of note, advanced *EGFR*-mutated NSCLCs treated with TKIs had worse outcomes in smokers than in non-smokers (30, 31). The transcriptomic activities of NSCLC driven by NSROs, including *EGFR*, in smokers, might account for these cancers’ higher resistance to standard TKIs and suggests the possibility of better responses to therapies such as chemotherapy or CDK4/6 inhibitors that target the cell cycle (65). This warrants further investigation regarding treatment selection for patients with NSRO-driven NSCLC.

In summary, based on multi-region whole-exome and RNA sequencing data, we have elucidated the clonal architectures and genomic features of three groups of East-Asian NSCLC: NSRO-driven in non-smokers, NSRO-driven in smokers, and typical-smoking tumors. We found no evidence that tobacco smoking affects clonal architecture or patterns of genomic alteration in NSRO-driven NSCLC. However, some transcriptomic pathway activities were more similar between NSRO-driven tumors in smokers and typical-smoking tumors than between NSRO-driven cancers in smokers and non-smokers. The in-depth analysis of NSRO-driven NSCLC in smokers and non-smokers presented here may provide guidance for optimizing treatment approaches.

## Methods

### Patients and clinical outcomes

Patients diagnosed with NSCLC at the National Cancer Centre Singapore (between 2013 and 2017) who underwent surgical resection of their tumors prior to receiving any form of anti-cancer therapy were enrolled in this study. Clinical information and histopathological features were obtained from the Lung Cancer Consortium Singapore. Written informed consent was obtained from all participants. This study was approved by the SingHealth Centralized Institutional Review Board (CIRB reference 2018/2963).

### Tumor/normal sample processing and whole-exome sequencing

Resected tumors and paired normal samples were sectioned and processed as previously described (24). Peripheral blood, or if peripheral blood was not available, normal lung tissue adjacent to the tumor was taken as a normal sample. The median number of sectors for an individual tumor was 3 (range 2-7, Supplementary Table S1). For whole exome sequencing, genomic DNA was extracted with the AllPrep DNA/RNA/miRNA Universal Kit (Qiagen), and 500 ng to 1 µg of genomic DNA was sheared using Covaris to a size of 300 to 400 bp. Libraries were prepared with NEBNext Ultra DNA Library Prep Kit for Illumina (New England Biolabs). Regions to sequence were selected with the SeqCap EZ Human Exome Library v3.0 (Roche Applied Science) according to the manufacturer’s instructions and underwent 2 × 151 base-pair sequencing on Hiseq 4000 (Illumina) sequencers. The median coverage of the capture target was 55.1X and 54.4X for normal and tumor samples, respectively (Supplementary Table S2).

### Somatic single nucleotide variation and insertion-deletion calling

Exome reads were trimmed with trimmomatic (version 0.39) to remove adaptor-containing or poor-quality sequences (66). Trimmed reads were mapped to the human reference sequence GRCh38.p7 (accession number GCA_000001405.22) using the BWA-mem software (version 0.7.15) with default parameters (67). Duplicate reads were marked and removed from variant calling using Sambamba (version 0.7.0) (68). Global mapping quality was evaluated by Qualimap 2 (version 2.2.1, Supplementary Table S2) (69). Somatic single nucleotide variations (SNVs) and insertion-deletions (indels) were called by MuTect2 (version 4.1.6.0) and Strelka2 (version 2.9.2) with default parameters (70, 71). We considered only variants called by both variant callers and with (i) ≥ 3 reads supporting the variant allele in the tumor sample, (ii) sequencing depth ≥ 20 in both the normal and tumor samples, and (iii) variant allele fraction ≥ 0.05. Variants were annotated by wANNOVAR (https://wannovar.wglab.org/) (72). Driver status of genes was based on the Catalog of Somatic Mutations in Cancer (COSMIC) database, downloaded February 24, 2021 (https://cancer.sanger.ac.uk/census) (73).

We excluded 12 out of 185 sectors (6.5%) that had tumor purity < 0.1 as assessed by HATCHet (74) (Supplementary Table S2). In tumor sectors with low tumor purity, fewer variants were called than in other sectors of the same tumor, supporting the estimation of low tumor purity (Supplementary Fig. S11).

### Definitions of truncal mutation, branch mutation, tumor mutational burden, and intra-tumor heterogeneity

We refer to mutations present in every sector of a tumor as “truncal”, and we refer to other mutations as “branch”. We defined tumor mutational burden (TMB) as the mean number of unique non-silent (nonsynonymous or splice-site) mutations across all sectors of a tumor. We defined intra-tumor heterogeneity (ITH) as the mean proportion of the number of unique branch mutations across all sectors. The clonality of somatic variants was evaluated by using MutationTimeR R package (75).

### Phylogenetic analysis

We used the Python PTI package (https://github.com/bioliyezhang/PTI, version 1.0) using the input of a “binary matrix” to infer phylogenetic relationships based on non-silent mutations (76).

### Mutational signature assignment and spectrum reconstruction

Mutational signature assignment was carried out with mSigAct R package (version 2.3.2, https://github.com/steverozen/mSigAct) and COSMIC mutational signature database (version 3.2) (77). For mutational signature analysis, we used all single-base substitution (SBS), including exonic and non-exonic variants. To better estimate the impact of smoking on cancer evolution, we first used the SignaturePresenceTest function with default parameters on all individual sectors within each group to decide whether the SBS4 mutational signature (the signature of tobacco smoking) was present in the sector’s mutational spectrum. In brief, SignaturePresenceTest estimates optimal coefficients for the reconstruction of the observed spectrum using the mutational signatures previously detected in NSCLC (40). The test does this without the SBS4 signature (null hypothesis) and with the SBS4 signature (alternative hypothesis). The test then carries out a standard likelihood ratio test on these two hypotheses to calculate a p-value. We then calculated Benjamini-Hochberg false discovery rates across all sectors of all tumors within the group.

To estimate the contribution of signatures to each spectrum we used the SparseAssignActivity function and the signatures found in lung adenocarcinomas in reference (40), except that SBS4 was included only if the false discovery rate based on the SignaturePresenceTest was < 0.05. We excluded SBS3 (caused by defective homologous recombination DNA damage repair mechanism) from sparse assignment after ensuring no pathogenic germline or somatic alterations in the *BRCA1* or *BRCA2* genes. Supplementary Table S7 and S8 show the SBS mutational spectra and signature activities of each sector. We did not assign activities for indels or double-base substitutions because there were too few of these mutations (median exome-wide counts of 10 and 1, respectively).

### Detection of fusion transcripts

We used STAR-Fusion (version 1.10.0) (78) to detect transcript fusions in the RNA-sequencing data with default parameters. We required candidate fusions to satisfy the following criteria:

- spanning fragment count ≥ 1
- junction read count + spanning fragment count ≥ 5
- presence of a large anchor-support read, and
- for intrachromosomal fusion partners, a genomic distance ≥ 1MB between fusion breakpoints

Supplementary Table S3 provides the full list of putative fusions.

### RNA sequencing and gene expression subtype

Total RNA was extracted and processed from 103 tumor samples as previously described (79). We used STAR (version 2.7.3a) to align raw RNA sequence reads to the human genome (GRCh38p7 build) and to estimate transcript abundance based on the reference transcriptome (GRCh38.85 build) (80). Only the counts of protein-coding genes were included for downstream analysis.

### Transcriptomic pathway analysis

Raw gene expression levels were transformed to transcript levels in transcripts per million (TPM) values (81). We computed pathway enrichment scores with the GSVA R package (version 1.40.1) and the Reactome subset of the Molecular Signatures Database (MSigDB, version 7.5.1) (43, 82, 83). Differential pathway expression was assessed using the limma R package (version 3.48.0) (84). Pathways with a Benjamini-Hochberg false discovery rate < 0.05 were taken as significant. Assignment of gene expression subtypes (terminal respiratory unit, TRU, versus non-TRU) was carried out as described (85). Gene expression values and pathway enrichment scores were transformed to Z-scores (mean of 0 and standard deviation of 1) before downstream analysis. Heatmaps were constructed with the ComplexHeatmap R package (version 2.8.0) (86). Heatmap columns were first clustered based on all rows using ComplexHeatmap::Heatmap function using default arguments for clustering distance and method, and then ordered by main group, patient, and gene expression status accordingly. Immune cell deconvolution was performed by using the CIBERSORT web application (https://cibersortx.stanford.edu/) (87).

## Data Availability

All WES and RNA sequencing data were deposited at the European Genome-phenome Archive (EGA, http://www.ebi.ac.uk/ega/), under the accession number EGAS00001006942. Supplementary Data and Code, including lists of somatic mutations, mutation-timing and clonality output from the MutationTimeR package, gene fusions, gene expression matrices, transcriptomic pathway activities, and immune cell deconvolution, are publicly available at https://github.com/Rozen-Lab/oncogene-NSCLC/.

## Authors’ Contributions

**C.-Y. Huang**: Data curation, formal analysis, visualization, methodology, writing original drafts, reviewing, and editing. **N. Jiang**: Formal analysis, methodology, writing, reviewing, and editing. **M. Shen**: Project coordination. **G. Lai**, **A.C. Tan**, **A. Jain**, **S.P. Saw**, **M.K. Ang**, **Q.S. Ng**, **D.W.T. Lim**, **R. Kanesvaran**, **E.H. Tan**, **W.L. Tan**, **B.H. Ong**, **K.L. Chua**, and **D. Anantham**: Clinical data curation and patient sample collection. **A. Takano** and **T.K.H. Lim**: Data curation, investigation, and methodology. **W.L. Tam**: Data curation and resources. **N.L. Sim**: Data curation, investigation, and resources. **A.J. Skanderup**: Conceptualization, data curation, resources, methodology, funding acquisition, writing, reviewing, and editing. **D.S.W. Tan**: Conceptualization, data curation, patient sample collection, resources, methodology, funding acquisition, supervision, writing, reviewing, and editing. **S.G. Rozen**: Conceptualization, supervision, resources, methodology, writing, reviewing, and editing.

## Acknowledgements

This work was funded by the National Medical Research Council (NMRC; Singapore) through the Translational and Clinical Research Program (NMRC/TCR/007-NCC/2013), Open Fund Large Collaboration Grant (NMRC/OFLCG/002/2018). This work was funded in part by grants from the Chang Gung Memorial Hospital (CGMH) Foundation (Grant No. CMRPG3F1911) to Chen-Yang Huang.

We thank Jacob J.S. Alvarez, and Jia Chi Yeo from the Genome Institute of Singapore, for data transfer. We thank Willie Yu from Duke-NUS Medical School, for helping with some Unix/Python scripts and setting up some bioinformatic pipelines. We thank Xinyi Yang from the Duke-NUS Medical School for testing the R codes. We thank the Lung Cancer Consortium Singapore (LCCS) for assisting with specimen collection and clinical-data compilation. Finally, we are grateful to the patients, physicians, and pathologists at the National Cancer Centre Singapore and Singapore General Hospital who contributed patient material.

