## Supplementary Figures S1-S11 for "Dearth of smoking-induced mutations in NSRO-driven non-small-cell lung cancer despite smoking exposure"

Affiliations
[1]Centre for Computational Biology, Duke-NUS Medical School, Singapore 169857, Singapore
[2]Programme in Cancer and Stem Cell Biology, Duke-NUS Medical School, Singapore 169857, Singapore
[3]Division of Hematology-Oncology, Department of Internal Medicine, Linkou Chang Gung Memorial Hospital and Chang Gung University, Taoyuan 333, Taiwan
[4]Division of Medical Oncology, National Cancer Centre Singapore, Singapore 168583, Singapore
[5]Institute of Molecular and Cell Biology, Agency for Science, Technology and Research (A*STAR), Singapore 138632, Singapore
[6]Department of Cardiothoracic Surgery, National Heart Centre Singapore, Singapore 169609, Singapore
[7]Division of Radiation Oncology, National Cancer Centre Singapore, Singapore 168583, Singapore
[8]Department of Respiratory and Critical Care Medicine, Singapore General Hospital, Singapore 169608, Singapore
[9]Department of Pathology, Singapore General Hospital, Singapore 169608, Singapore
[10]Genome Institute of Singapore, Singapore 138672, Singapore
[11]Cancer Science Institute of Singapore, National University of Singapore, Singapore 117599, Singapore
[12]Department of Biochemistry, Yong Loo Lin School of Medicine, National University of Singapore, Singapore 117597, Singapore
[13]Duke-NUS Medical School Singapore, Singapore 169857, Singapore
[14]Cancer Therapeutics Research Laboratory, Division of Medical Sciences, National Cancer Centre Singapore, Singapore 168583, Singapore
[15]NUS Graduate School for Integrative Sciences and Engineering, Singapore 117456, Singapore

*To whom correspondence may be addressed:

Anders J. Skanderup, PhD
Genome Institute of Singapore, Singapore
60 Biopolis St, Singapore 138672


Daniel S.W. Tan, MD, PhD
Division of Medical Oncology, National Cancer Centre Singapore
30 Hospital Boulevard, Singapore 168583


Steven G Rozen, PhD
Centre for Computational Biology, Duke-NUS Medical School, Singapore
8 College Rd, Singapore 169857


### Supplementary figure S1

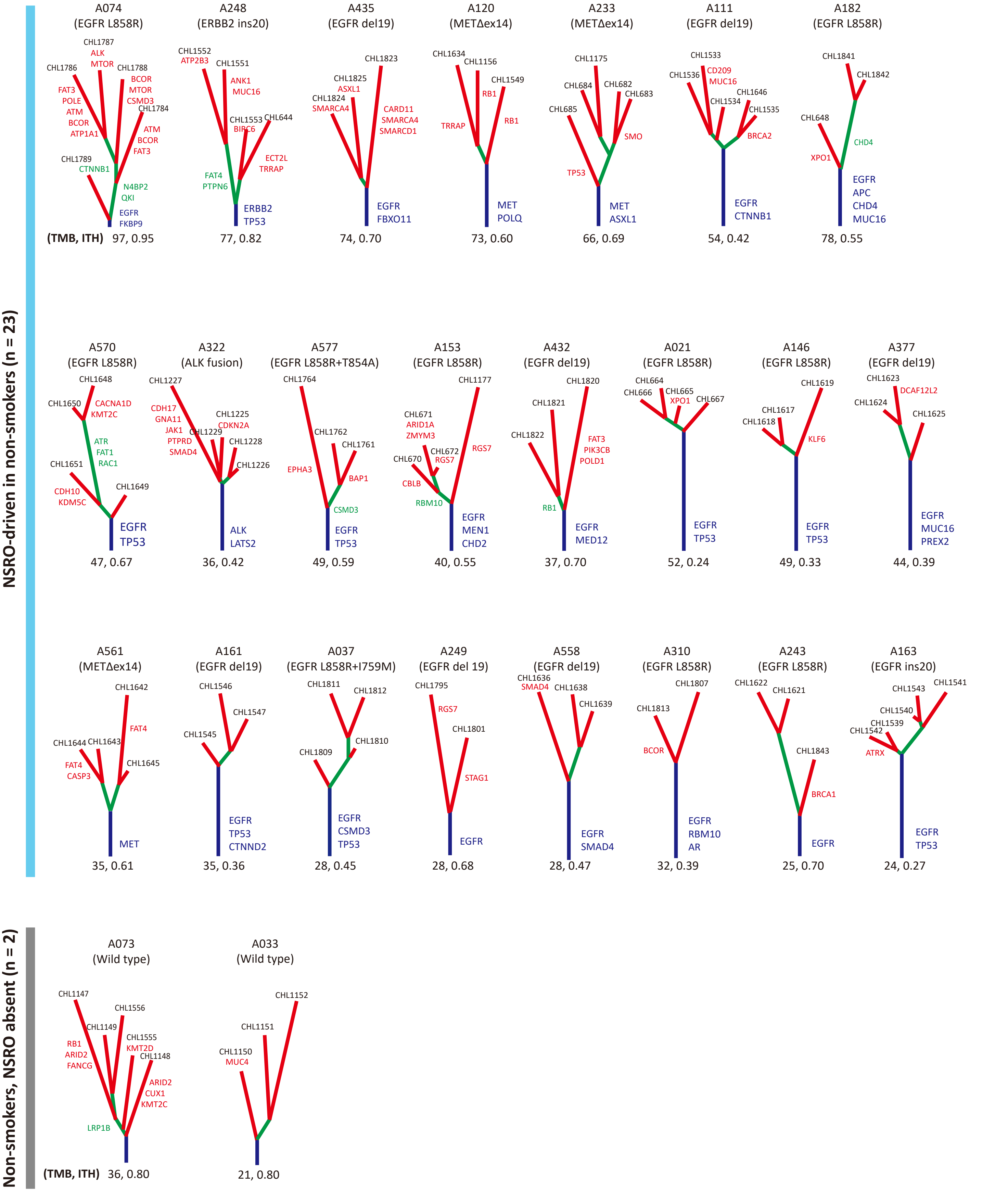

### Supplementary figure S1 (continued)

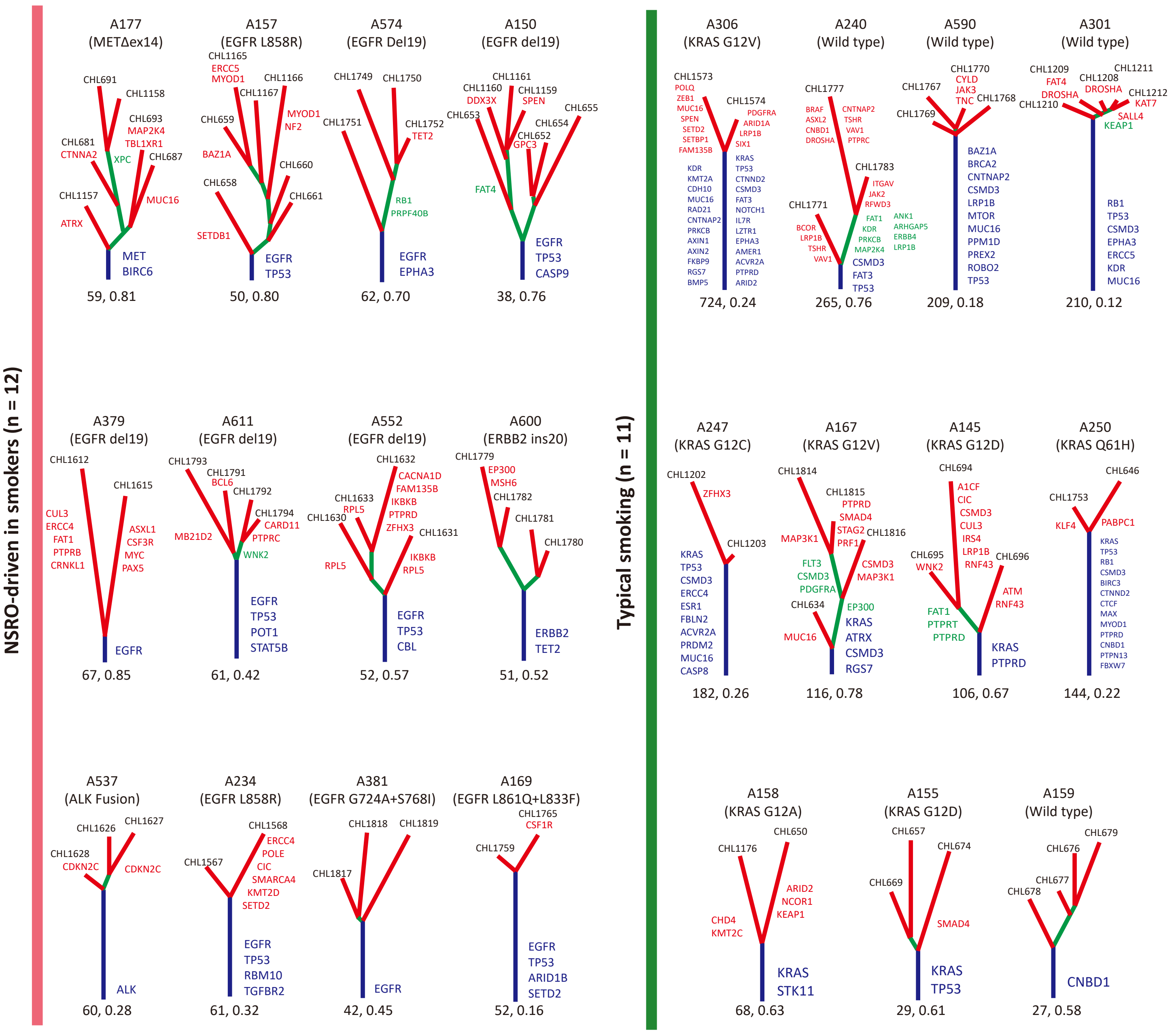


**Supplementary figure S1.** Phylogenetic trees of all tumors in the study. The tumor ID and key driver mutations are indicated above each tree. Branch tips are labeled with the corresponding sector IDs, as in Supplementary Table S2. Additional mutations in driver genes are shown along the branches. Within each tree, branch lengths are proportional to the total number of mutations along a given branch, but the scales are different across trees. Tumor mutational burden and intra-tumor heterogeneity are shown below each tree. See main text for methods for inferring phylogenies.

#### Supplementary figure S2

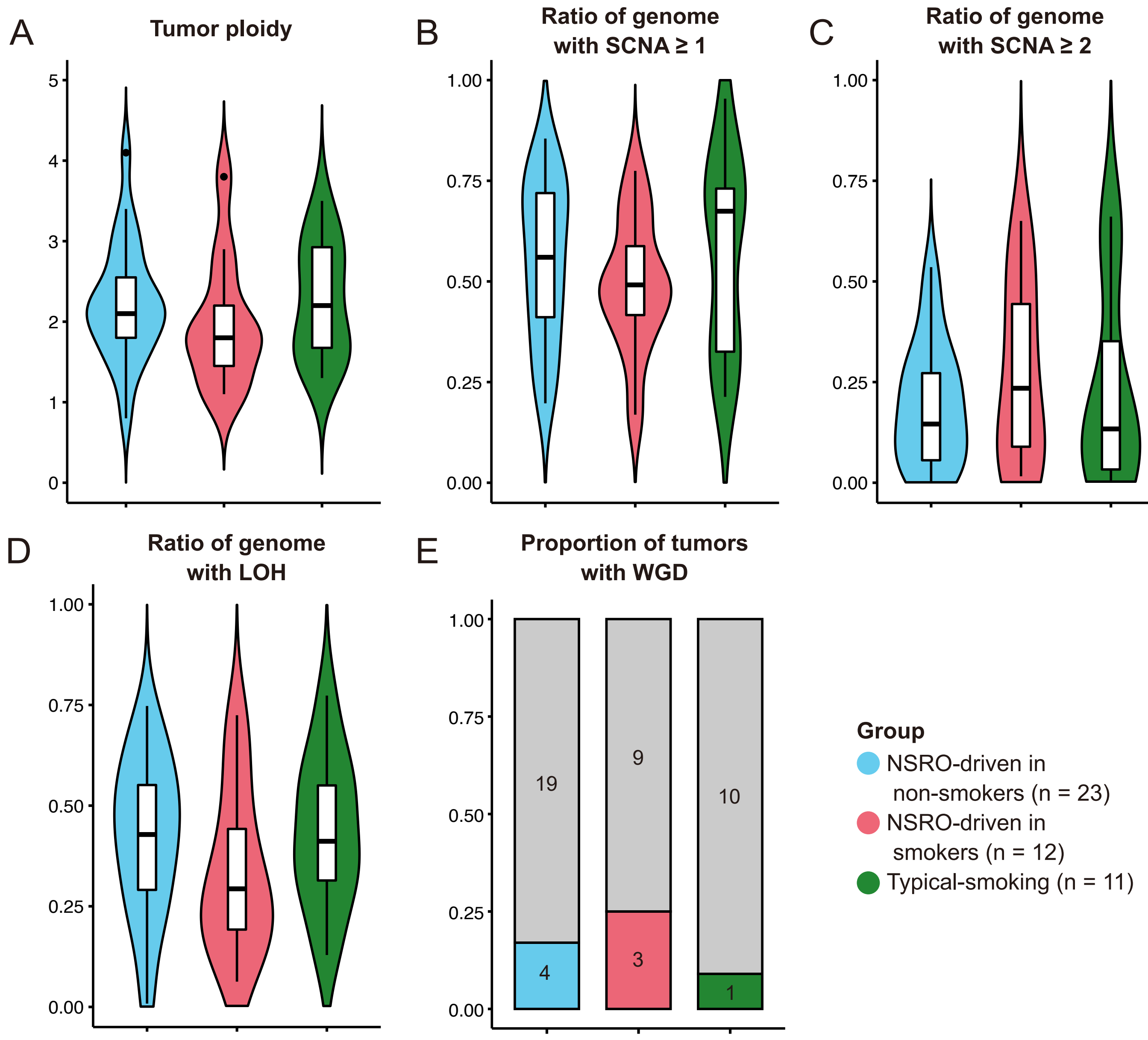


**Supplementary figure S2.** Comparison of the three tumor groups with respect to five parameters as indicated. SCNA: somatic copy-number alteration, WGD: whole-genome doubling.

To detect SCNA and WGD, for each tumor, we applied the HATCHet package (version 0.4.8, http://compbio.cs.brown.edu/hatchet/) to the mapped reads in all sectors from the tumor (Zaccaria and Raphael, 2020) (1). Germline single nucleotide polymorphisms (SNPs) were called using freebayes on the normal sample (version 1.3.0, https://github.com/freebayes/freebayes) (Garrison and Marth, 2012) (2). We considered SNPs with allele fraction between 0.4 and 0.6 and read depth > 20 to be heterozygous and therefore suitable as input for SCNA analysis. HATCHet estimated the tumor ploidy, purity, WGD status, and copy numbers of all the sectors of each tumor. We choose the two-clone solution (one normal clone and one tumor clone) inferred by HATCHet. The SCNA was calculated as the somatic copy number gain or loss relative to background ploidy, which was 4 if WGD was present, or 2 otherwise.

### Supplementary figure S3

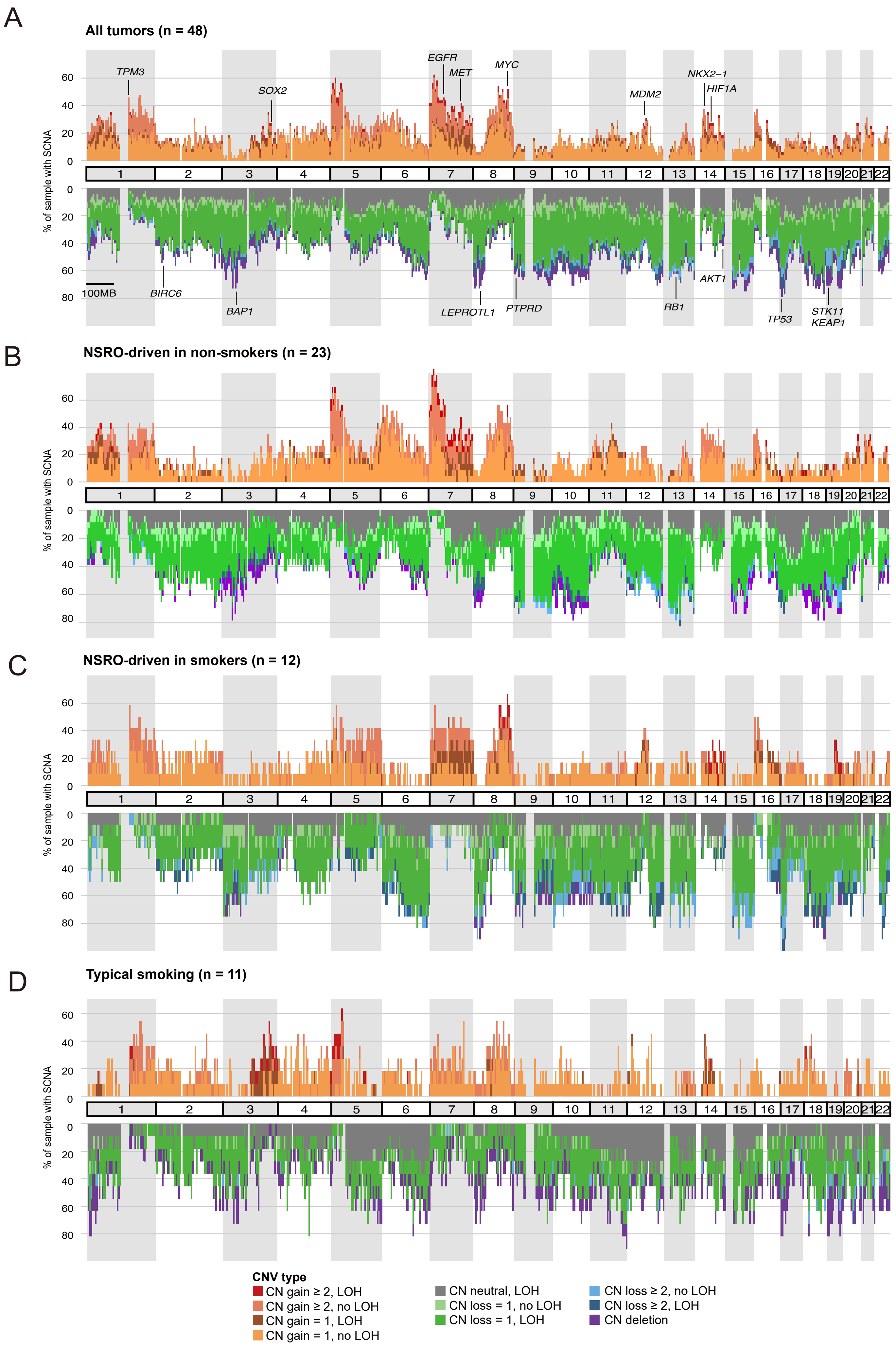

**Supplementary figure S3 (continued).** Genome-wide somatic copy number alterations in (A) all tumors in the study cohort (n = 48), (B) NSRO-driven in non-smokers (n = 23), (C) NSRO-driven in smokers (n = 12), and (D) Typical-smoking-related (n = 11).

Plotting code is at https://github.com/Rozen-Lab/oncogene-NSCLC/), and was based on HATCHet's allele-specific copy-number estimates for each tumor.

#### Supplementary figure S4

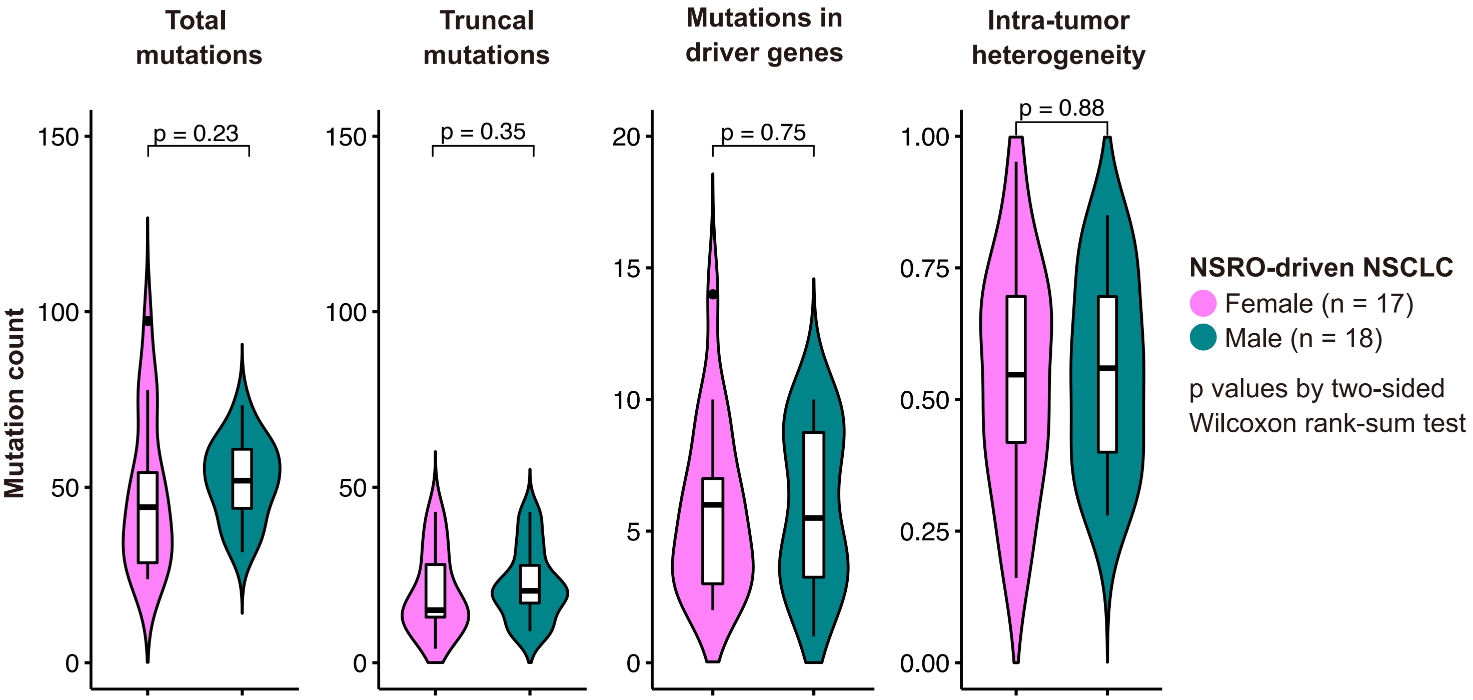


**Supplementary figure S4.** Counts of total mutations, truncal mutations, driver mutations, and levels of intra-tumor heterogeneity between oncogene-driven tumors in female and male patients.

### Supplementary figure S5

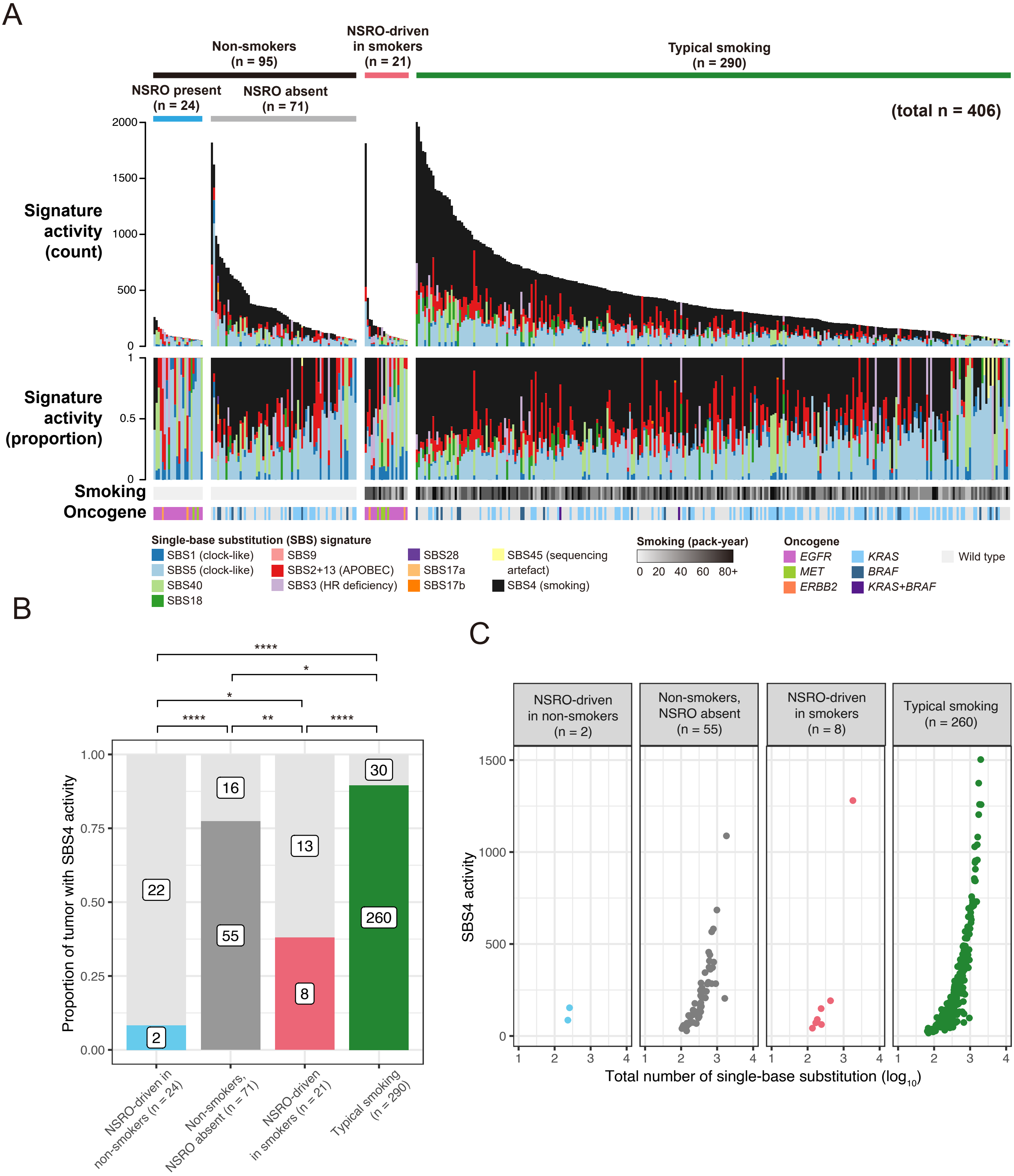


Two-sided Fisher's exact test, with
Benjamini-Hochberg correction,
q value: * < 0.1, ** < 0.01, *** < 0.001, **** < 0.0001

**Supplementary figure S5.** Analysis of single-base substitution (SBS) mutational signature activities in 406 Western lung adenocarcinomas (The Cancer Genome Atlas Research Network, 2014) (3). (A) Mutational-signature activity by absolute mutation counts (top panel) and proportion (second panel). Smoking status and presence of mutations in selected oncogenes are indicated below. (B) The proportion of tumors with SBS4 activity by tumor group. Q values by two-sided Fisher's exact tests with Benjamini-Hochberg correction. (C) SBS4 mutational activity in tumors showing SBS4 activity.

Supplementary figure S6

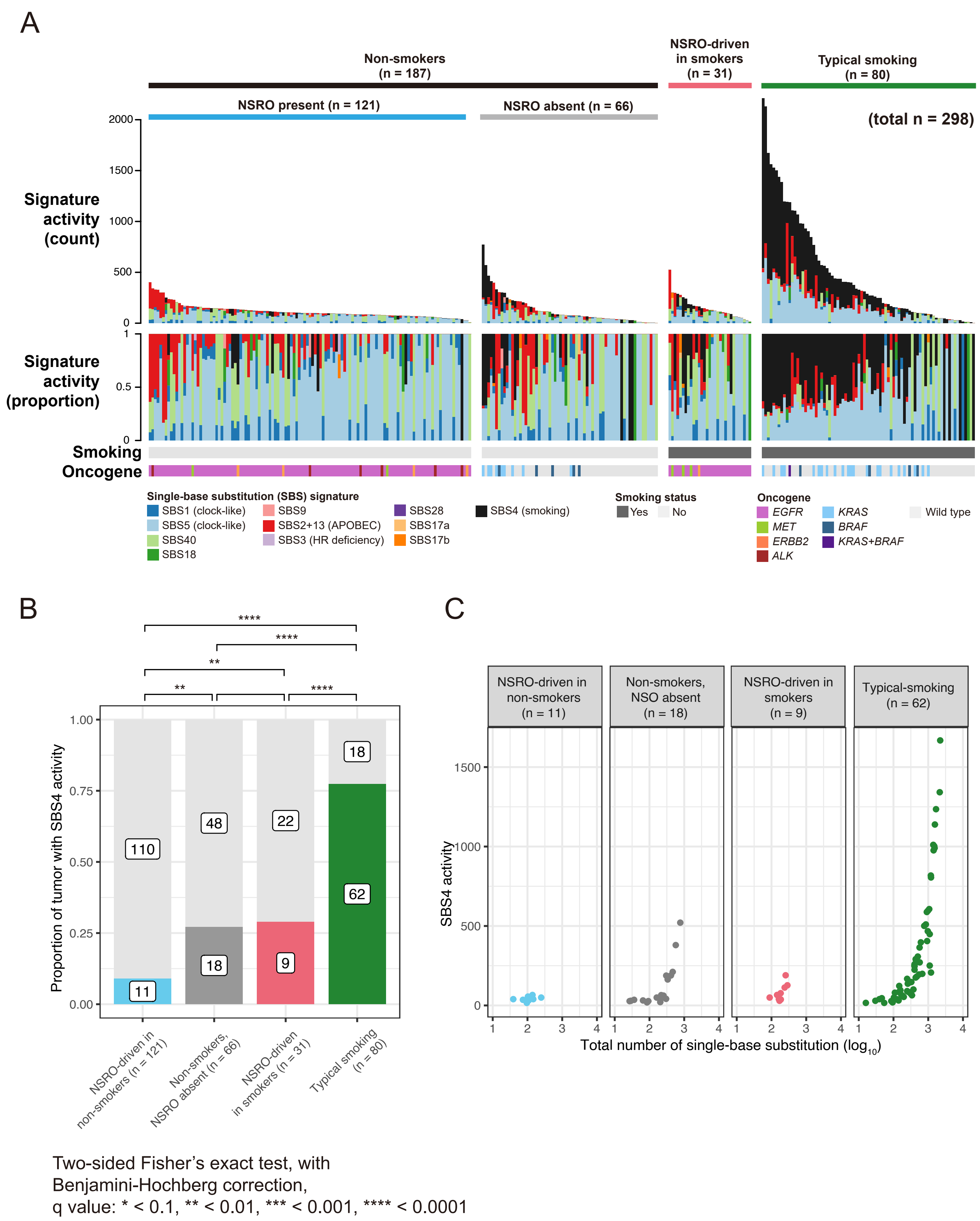


**Supplementary figure S6.** Analysis of single-base substitution (SBS) mutational signature activities in 298 East Asian lung adenocarcinomas (4). Panels A, B, C are as in Supplementary Fig. S5.

### Supplementary figure S7

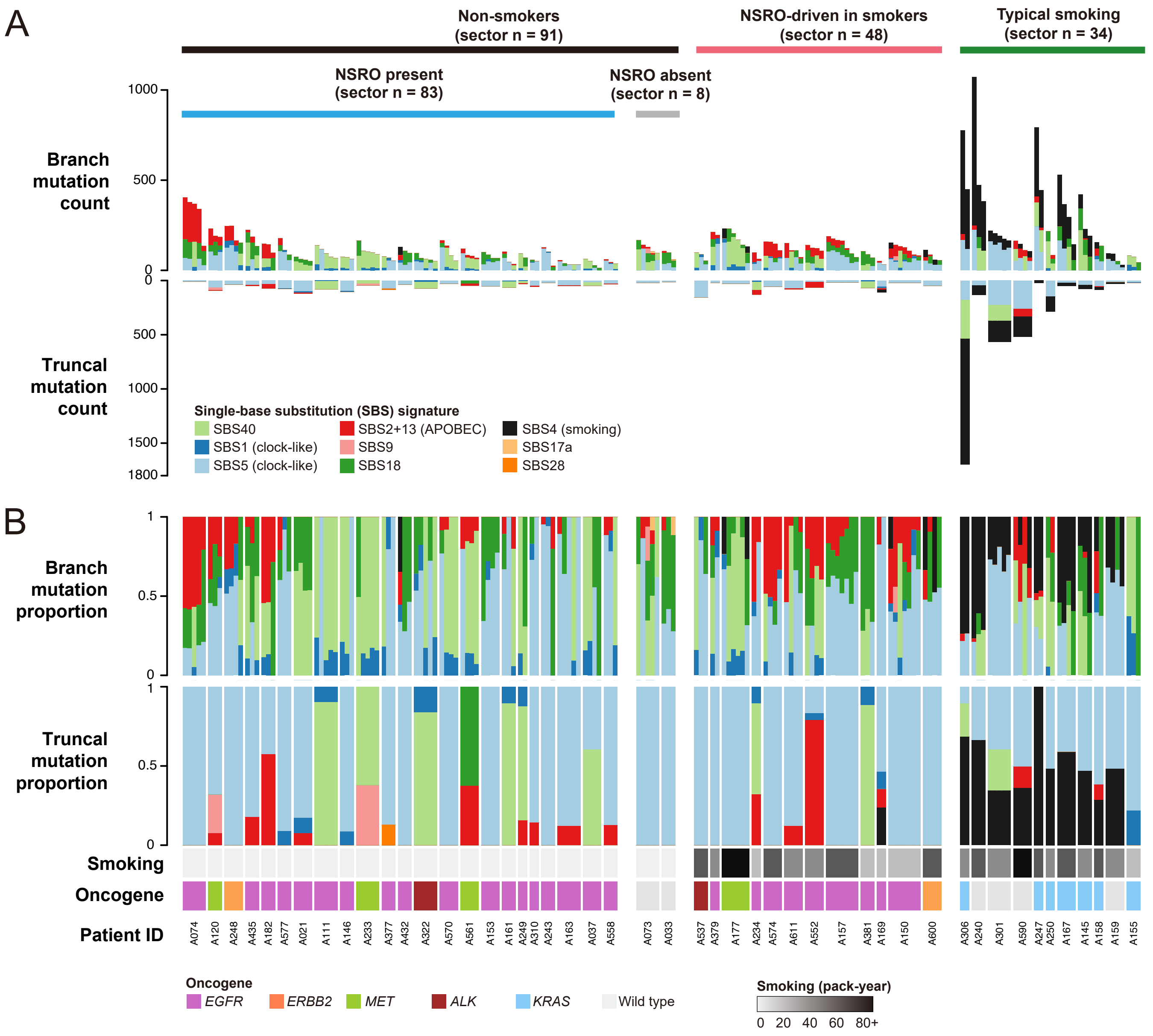


**Supplementary figure S7.** Comparison of the activities of single-base substitution (SBS) mutational signatures in branch versus trunk across the three groups. (A) Absolute counts and (B) proportions of mutations due to different mutational signatures in branches (upper panel) and trunks (lower panel). Smoking status and presence of oncogenes are indicated below. Counts and proportions are based on separate assignments of mutational signatures to truncal and branch mutations.

Supplementary figure S7 (continued)

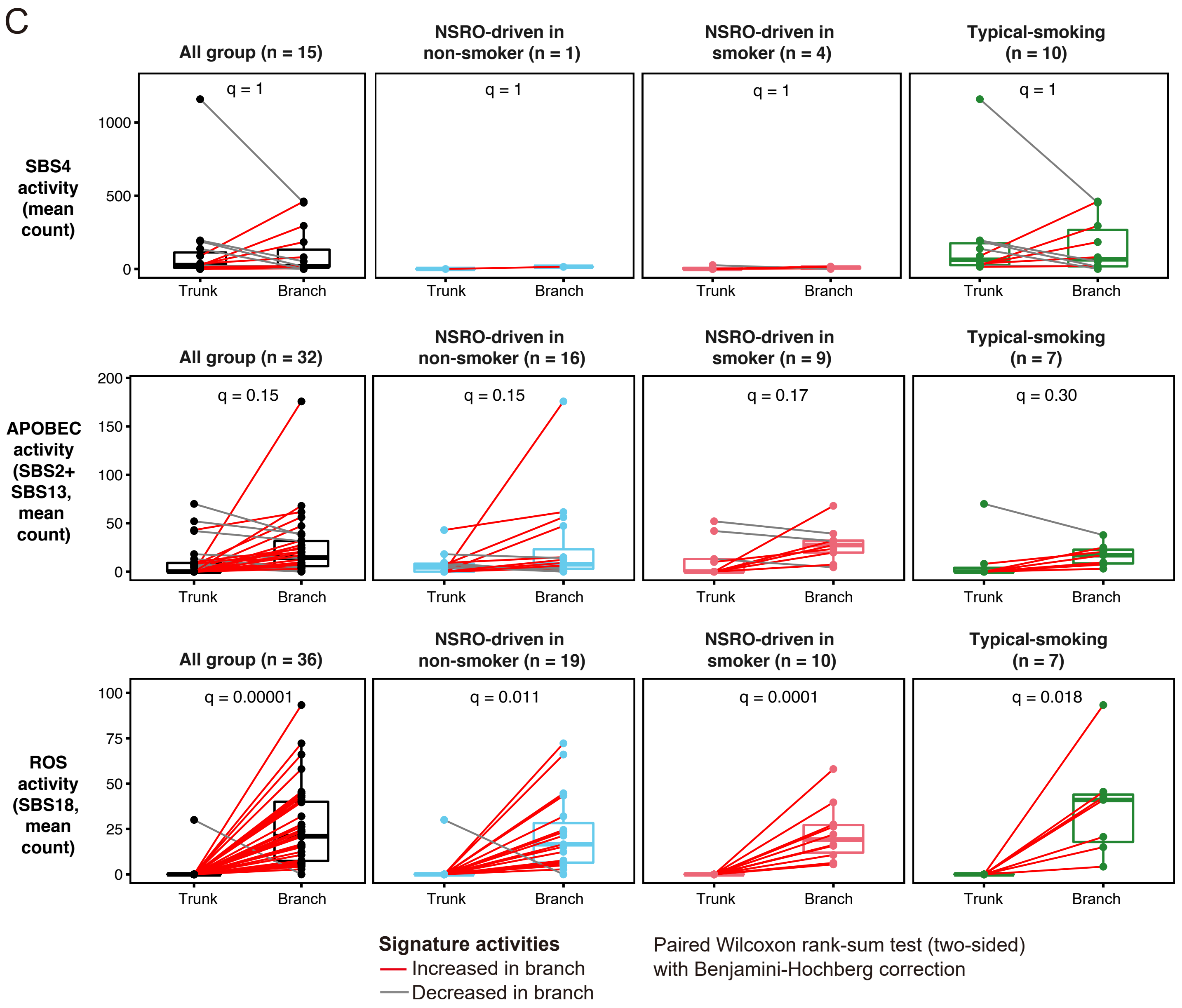


**Supplementary figure S7.** (continued) (C) Activities of 3 signatures in trunks versus branches. The y-axis position of each point represents mutations counts for the trunk or the branch of 1 tumor. Q-values by two-sided paired Wilcoxon rank-sum tests with Benjamini-Hochberg correction. ROS: reactive oxygen species.

### Supplementary figure S8

A

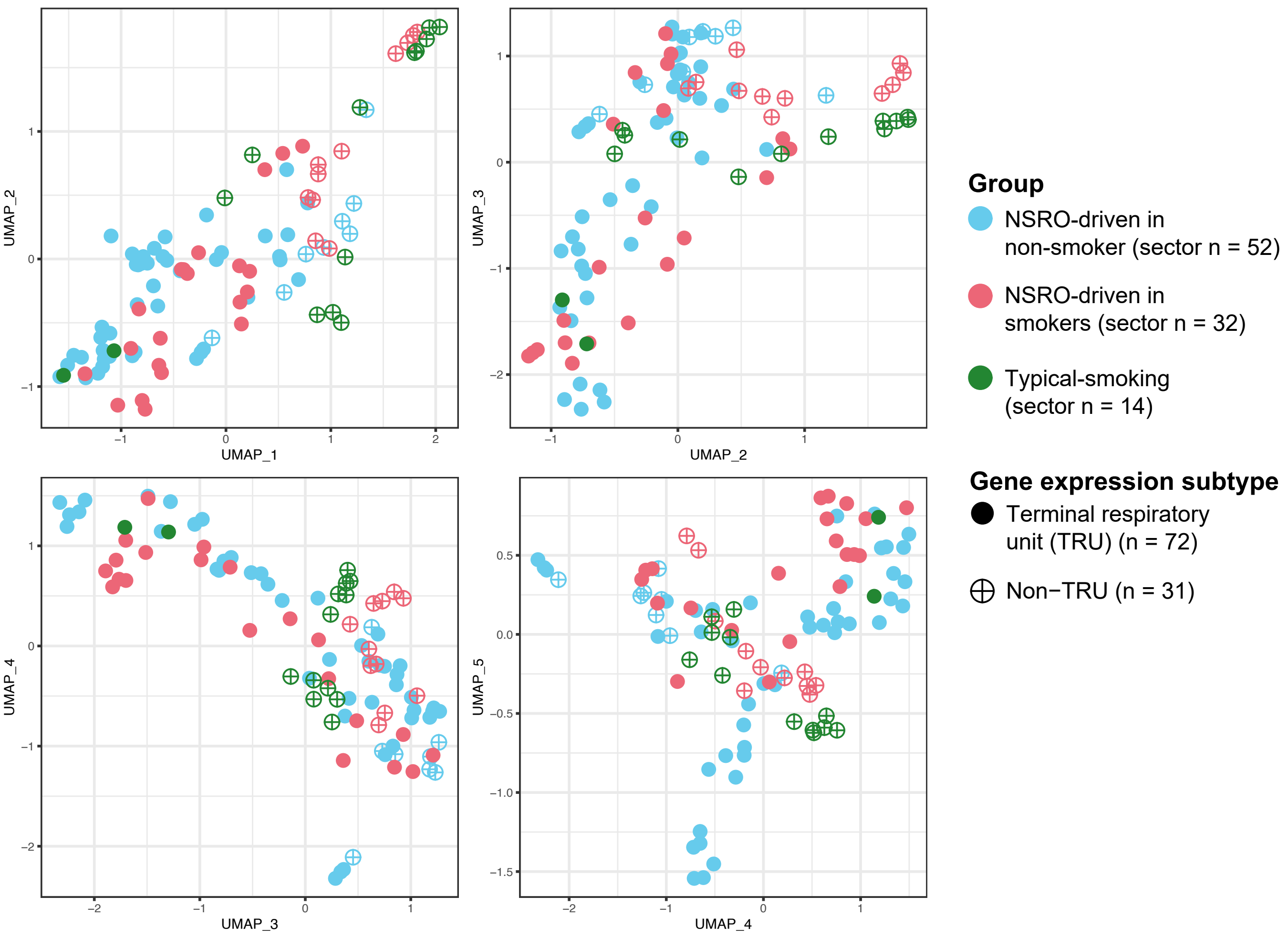


B

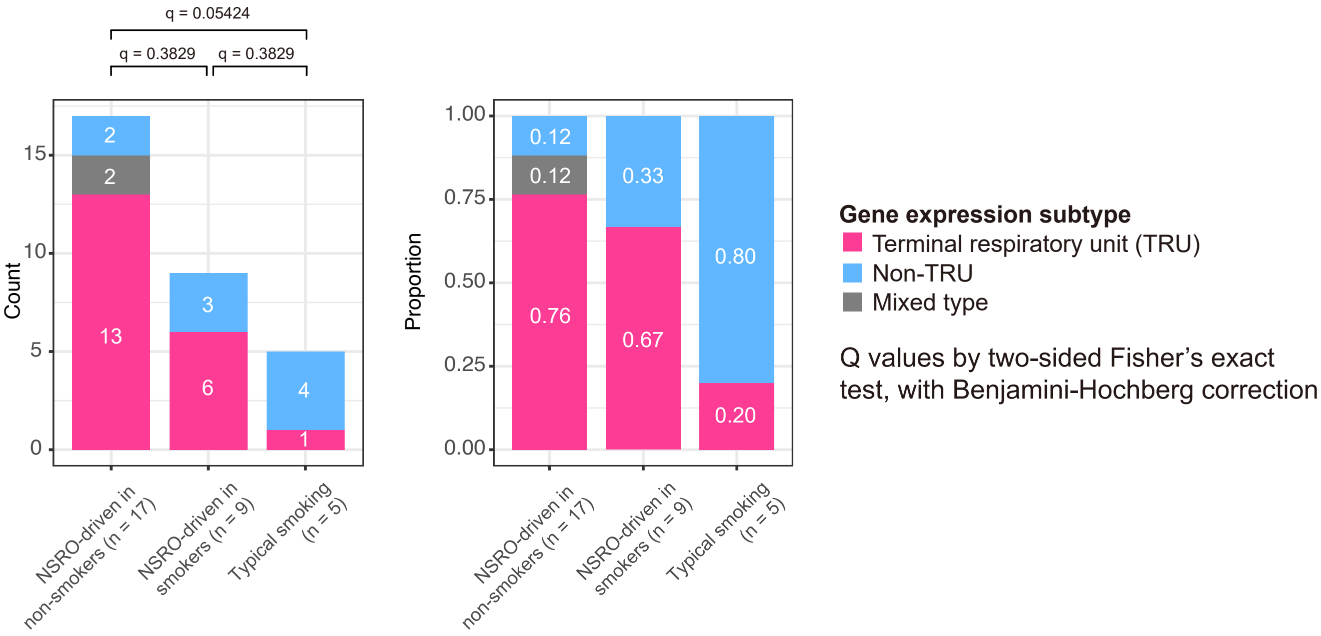


**Supplementary figure S8.** (A) UMAP projections of transcript levels of all coding genes by sector. TRU: terminal respiratory unit. (5) (B) Counts and proportions of TRU versus non-TRU gene expression-subtype tumors. Tumors with both TRU and non-TRU sectors were classified as 'mixed type'.

### Supplementary figure S9

A

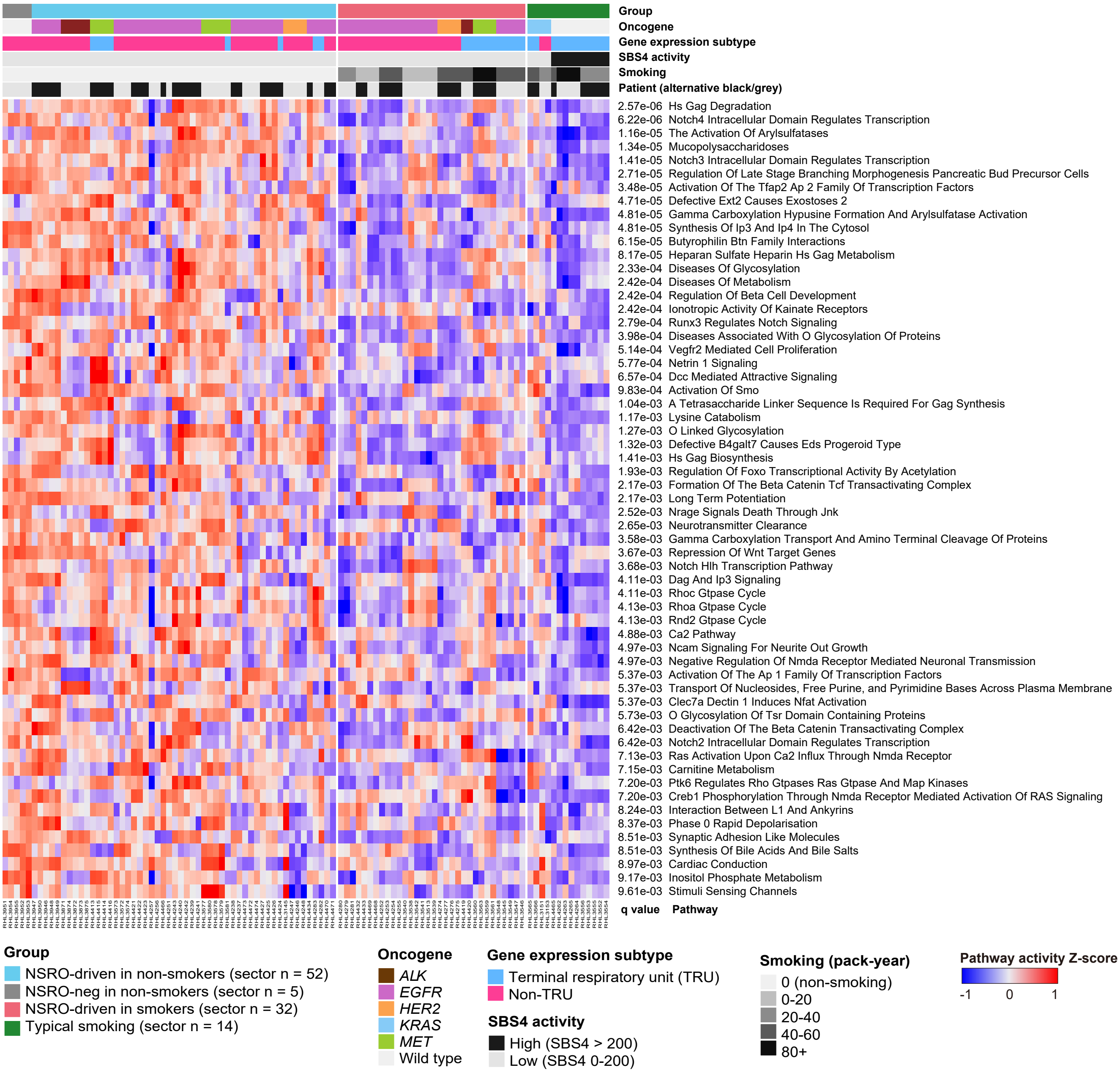


**Supplementary figure S9.** (A) Reactome pathways that were up-regulated in tumors in non-smokers versus smokers (6). Columns represent sectors and are grouped by tumor. Rows represent pathways. Pathways with a Benjamini-Hochberg false discovery rate < 0.01 are shown.

### Supplementary figure S9 (continued)

B

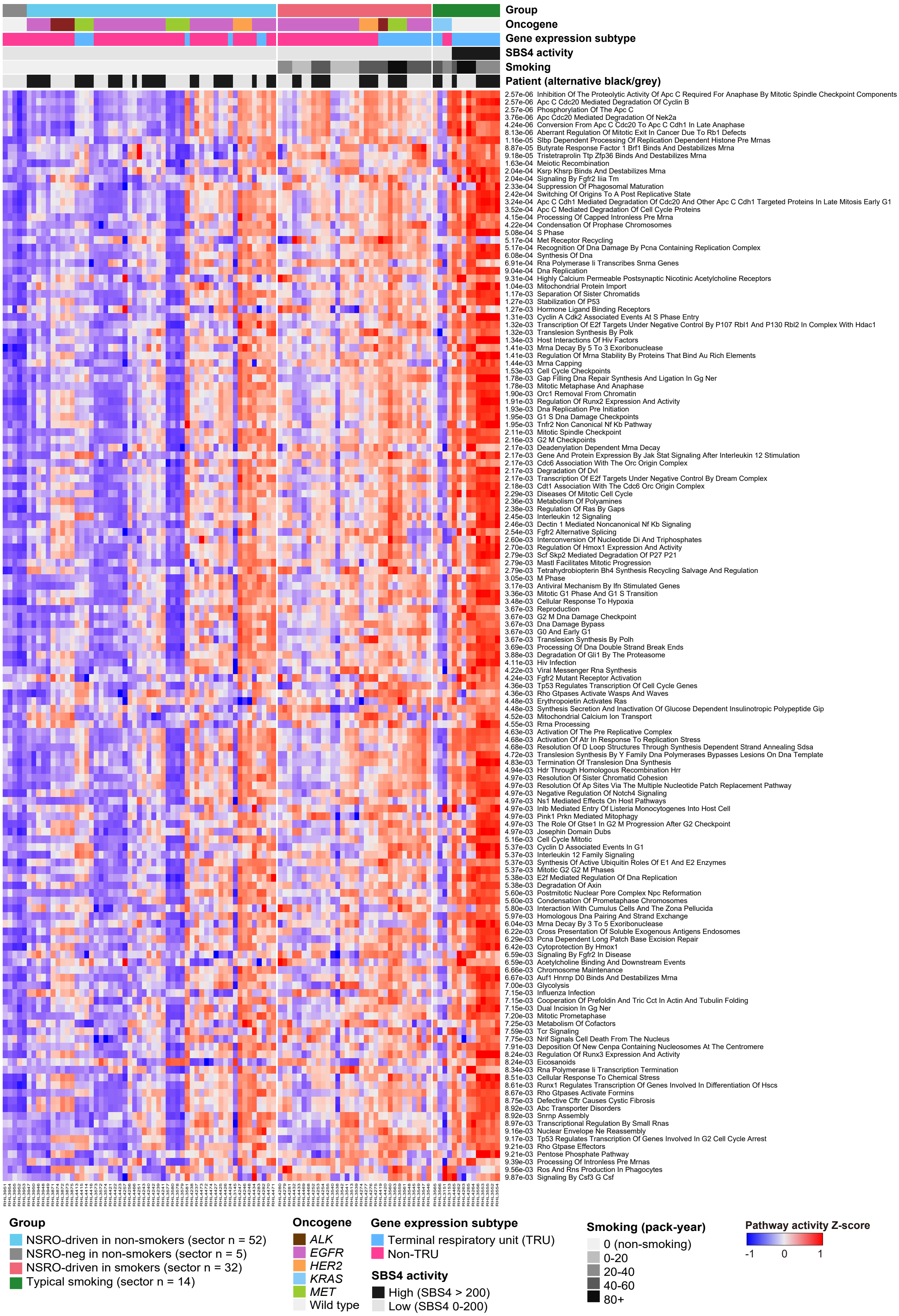


**Supplementary figure S9.** (continued) (B) Analogous to panels be for pathways down-regulated in tumors in non-smokers versus smokers.

### Supplementary figure S10

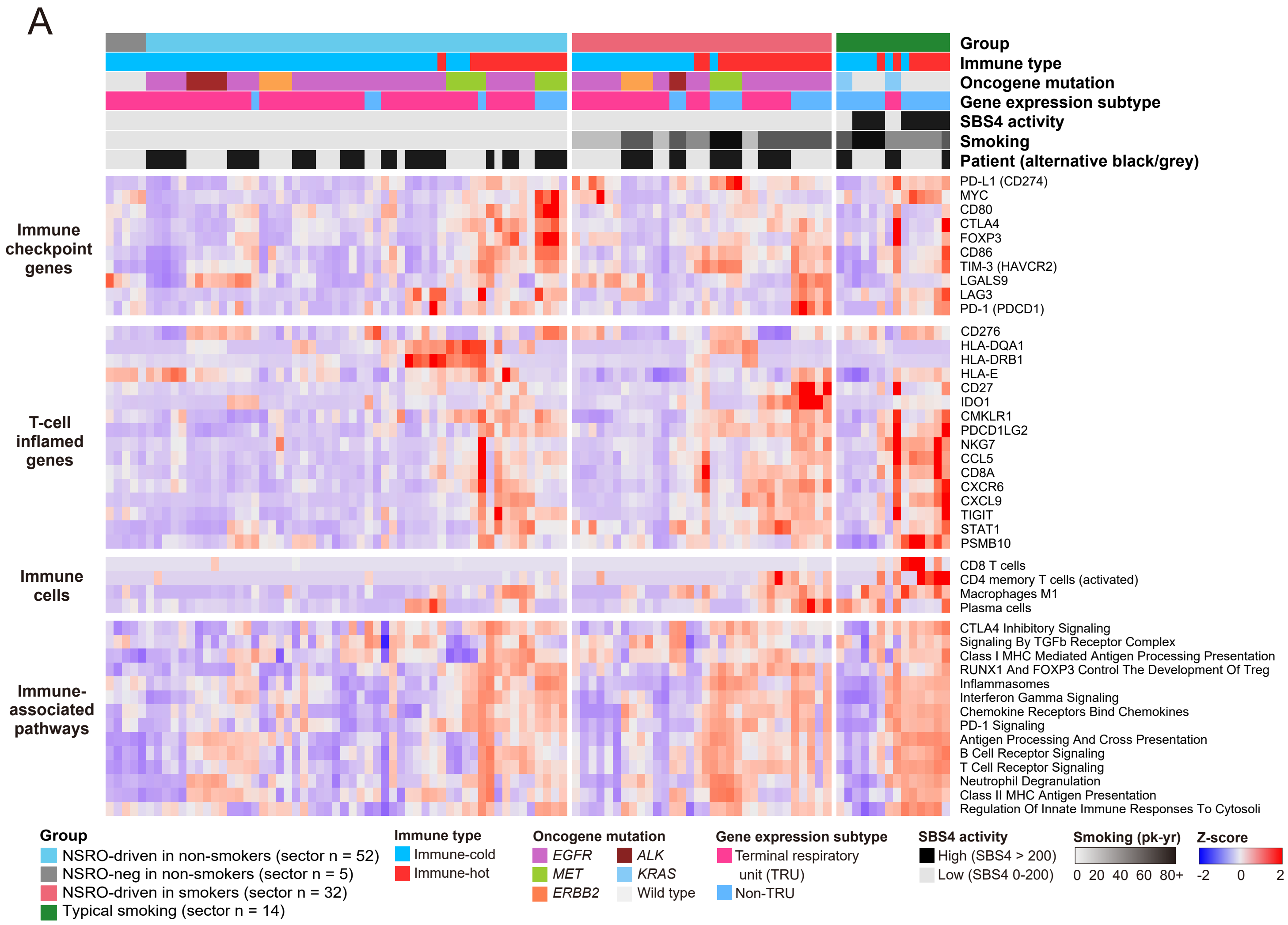


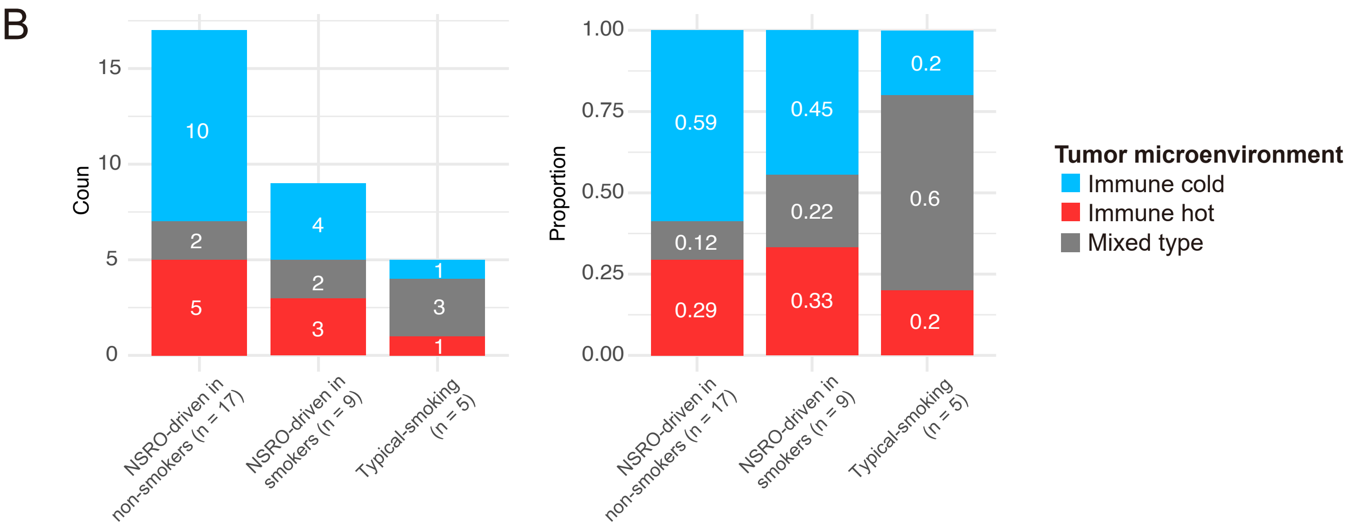


**Supplementary figure S10.** (A) Analysis of immune tumor microenvironment (TME) based on transcript levels of immune-checkpoint genes and T-cell-inflamed genes, plus proportions of CIBERSORT-estimated immune cells and activities of selected Reactome immune-associated pathways. (B) Proportion of immune-hot versus immune-cold (see Methods) TMEs in three tumor groups. Tumors with both immune-hot and immune-cold sectors are classified as 'mixed type'.

### Supplementary figure S11

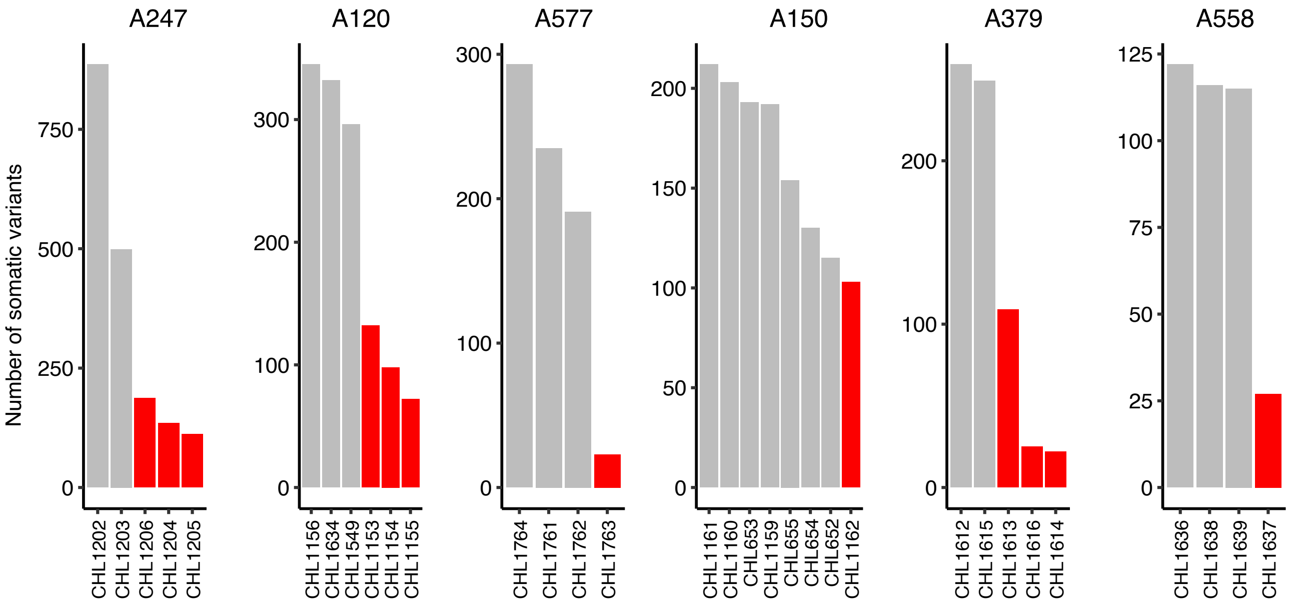


**Supplementary figure S11.** Numbers of somatic variants by sector for tumors with sectors with low purity. Tumor sectors with tumor purity < 0.1 (shown in red) were excluded from analysis because many somatic mutations in these sectors were likely not detected.
